# Cell-surface receptor-mediated regulation of synaptic organelle distribution controls dendritic spine maturation

**DOI:** 10.1101/2024.05.08.592949

**Authors:** Ben Verpoort, Luísa Amado, Jeroen Vandensteen, Elke Leysen, Dan Dascenco, Joris Vandenbempt, Irma Lemmens, Joris Wauman, Kristel Vennekens, Abril Escamilla-Ayala, Ana Cristina Nogueira Freitas, Thomas Voets, Sebastian Munck, Jan Tavernier, Joris de Wit

## Abstract

The spine apparatus (SA), an endoplasmic reticulum-related organelle present in a subset of mature dendritic spines, plays a key role in postsynaptic development and has been implicated in various neurological disorders. However, the molecular mechanisms that dictate SA localization at selected synapses remain elusive. Here, we identify a postsynaptic signaling complex comprising the GPCR-like receptor GPR158 and a largely uncharacterized phospholipase C (PLC), PLCXD2, that controls SA abundance. Sparse genetic manipulations *in vivo* demonstrate that in the absence of GPR158, unrestrained PLCXD2 activity impedes postsynaptic SA incorporation and hampers dendritic spine maturation. Finally, we show that extracellular heparan sulfate proteoglycan (HSPG) binding modulates the GPR158-PLCXD2 interaction. Together, our findings reveal how a postsynaptic receptor signaling complex regulates the local lipid microenvironment to control SA abundance required for the proper maturation of dendritic spines.

## Introduction

Synapses, the specialized junctions connecting pre- and postsynaptic neurons into neural circuits, are molecularly and morphologically diverse, reflecting their distinct functional properties. This diversity plays a key role in circuit connectivity, plasticity and vulnerability in disease.^1–3^ A less appreciated form of synapse diversity is apparent at the organelle level. A striking example is the SA organelle, which is found in a small fraction of dendritic spines. This unique organelle consists of closely packed stacks of smooth endoplasmic reticulum (sER) cisternae that are connected to the tubular ER network in dendritic shafts.^4^ The SA preferably associates with large, mature dendritic spines^5–7^ and regulates various forms of synaptic plasticity in both rodent^8–11^ and human^12^ neurons, as well as the long-term stability of dendritic spines.^13^ Accordingly, inadequate control over SA localization and function has been implicated in several brain disorders.^14–16^ However, the mechanisms that determine how the SA is recruited to and stabilized in specific dendritic spines remain elusive.

The sER dynamically enters and exits dendritic spines, visits that are typically transient.^17^ Only in dendritic spines where the sER has been stably inserted on a scale of hours will a SA develop.^17^ Myosin Va (myoVa), an F-actin-based motor protein, in conjunction with caldendrin, increases sER dwell time and facilitates SA formation.^18,19^ In addition, interactions between ER-resident proteins and cell-surface proteins localized at the PM have been suggested to promote sER anchoring in dendritic spines. This is based on the observation that inhibiting proteolytic activity in cultured neurons and thus preventing the cleavage of numerous transmembrane proteins drastically increases spine sER content.^20^ ER-PM contact sites are present in various cell types, including neurons, and play a crucial role in regulating intracellular Ca^2+^ dynamics.^21^ In dendritic spines, these contact sites are established between tubular extensions emanating from the SA and the postsynaptic PM, and are observed in almost all SA-containing spines.^4,5,22–24^ Tethering proteins that maintain the nanoscale apposition between both compartments constitute key components of ER-PM contact sites and are predominantly secured by protein-lipid interactions between integral ER proteins and negatively charged PM phosphoinositides, including phosphatidylinositol 4,5-bisphosphate (PIP2).^25^ Cell-surface protein-mediated interactions with the ER, either by direct binding to ER resident proteins or indirectly via a modification of the local membrane lipid microenvironment, would constitute an attractive, spatiotemporally precise way to govern the specific localization of the SA at a subset of dendritic spines.

We previously identified GPR158, a postsynaptic GPCR-like cell-surface protein, as an input-specific organizer that controls the development of hippocampal mossy fiber- CA3 synapses.^26^ GPR158 is an unconventional member of the family of class C GPCRs associated with mood disorders^27^ and cognition.^28^ Its extracellular binding partners and associated signaling pathways have only recently begun to be elucidated. Extracellularly, GPR158 interacts with presynaptic HSPGs^26,29^ and acts as a metabotropic glycine receptor.^30^ Intracellularly, GPR158 signals non-canonically via its association with the regulator of G protein signaling 7 (RGS7)/Gβ5 complex.^31,32^ Whether GPR158 operates solely through the RGS7/Gβ5 complex across various neural circuits or whether additional, context-dependent signaling mechanisms exist is not known.

Here, using an unbiased protein-protein interaction screen, we identify PLCXD2, an atypical PLC with no known function in the brain, as a novel intracellular GPR158 interactor. We find that PLCXD2 is a constitutively active PLC that associates with ER-PM contact sites to limit their formation and function. Binding of GPR158 to PLCXD2 prevents continuous PIP2 depletion and restores the formation of functional ER-PM contact sites. PLCXD2 localizes to dendritic spines where it hinders the incorporation of the SA organelle. Sparse conditional knockout (cKO) of *Gpr158* in cortical neurons *in vivo* leads to a loss of SA-bearing spines accompanied by a reduction in the fraction of mature dendritic spines. In agreement with GPR158’s inhibitory control over PLCXD2, removal of *Plcxd2* in *Gpr158* cKO neurons rescues SA content and dendritic spine maturation. Finally, we demonstrate that HSPG-binding modulates the GPR158- PLCXD2 complex. Together, our findings reveal how a novel GPR158-PLCXD2 signaling complex regulates the local lipid microenvironment to control postsynaptic SA content required for the proper maturation of dendritic spines.

## Results

### Identification of a novel postsynaptic GPR158-PLCXD2 complex

To gain more insight into the signaling mechanisms employed by GPR158 we conducted an unbiased, high-throughput two-hybrid screen termed mammalian protein-protein interaction trap (MAPPIT).^33^ Using the cytoplasmic domain of GPR158 as bait, we tested a library of approximately 15.000 prey proteins from the human ORFeome collection for functional complementation of a signaling-deficient chimeric cytokine receptor **(Figure 1A)**. To prioritize the hits resulting from this screen, we focused on potential interactors that are co-expressed with *Gpr158*. Layer 4 (L4) neurons in the barrel field of the developing primary somatosensory cortex show a striking enrichment of *Gpr158* expression, as visualized by β-Galactosidase immunohistochemistry on coronal brain sections from *Gpr158^LacZ/+^* mice **(Figure S1A)**. We cross-referenced the list of putative GPR158 interactors with L4-enriched transcripts gathered from 3 publicly available barrel cortex transcriptome datasets.^34–36^ **(Figure S1B)**. Next, co-expression of *Gpr158* and L4-specific putative interactors was assessed using the Mouse Brain Atlas (http://mousebrain.org).^37^ Based on its striking co-expression with *Gpr158* in select populations of neurons, we selected phospholipase C X-domain containing 2 (PLCXD2), an atypical PLC with no known function in the brain, for further validation.

**Figure 1.**
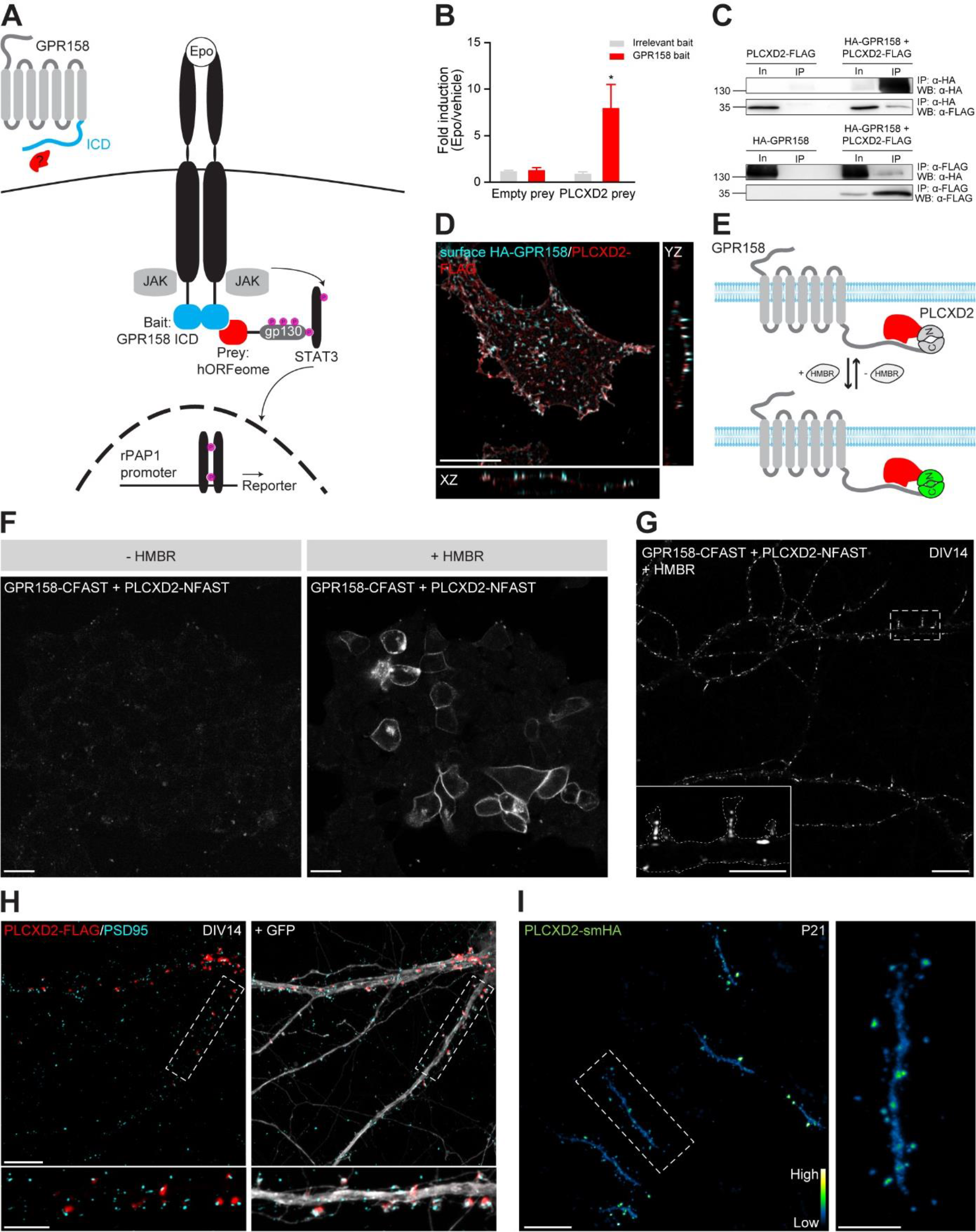
Identification of a novel postsynaptic GPR158-PLCXD2 complex (see also Supplementary figures S1 and S2). **(A)** Schematic representation of MAPPIT. The bait receptor consisted of GPR158’s cytoplasmic domain (ICD), excluding intracellular loops, fused to a signaling-deficient chimeric erythropoietin (epo)-leptin (lep) receptor (EpoR-LepR-F3). Prey proteins are tethered to a gp130 fragment containing functional STAT3 recruitment sites. Following epo stimulation, association of bait and prey proteins restores a functional complex resulting in STAT3 activation, whose activity can be monitored by a STAT3-responsive reporter. **(B)** Binary MAPPIT experiments confirm a specific interaction between PLCXD2 and GPR158’s cytoplasmic domain. EpoR-LepR-F3-GPR158 ICD and EpoR-LepR-F3- eDHFR (irrelevant bait) were introduced in HEK239T cells together with gp130- PLCXD2 or gp130 (empty prey). Bar graphs represent the fold induction from cells assayed in triplicate, defined as the ratio between the average luciferase activities of the epo-treated and untreated cells. \**P*<0.05, two-tailed unpaired t-test. **(C)** HA-GPR158 and PLCXD2-FLAG were expressed alone or together in HEK239T cells for 48 hours. Lysates were prepared, immunoprecipitated and blotted for HA or FLAG. **(D)** HEK293T cells expressing HA-GPR158 and PLCXD2-FLAG were live labeled for HA (cyan), then fixed, permeabilized and immunostained for FLAG (red). Orthogonal views show clear colocalization at the PM. Scale bar 10 µm. **(E)** Schematic representation of splitFAST. Following association between GPR158 and PLCXD2, the individual FAST fragments complement and fluoresce in the presence of HMBR. **(F)** HEK239T cells co-expressing GPR158-CFAST and PLCXD2-NFAST were visualized live with or without the addition of HMBR. Scale bar 20 µm. **(G)** DIV14 cultured mouse cortical neurons expressing GPR158-CFAST and PLCXD2- NFAST were visualized live in the presence of HMBR. Scale bar 10 µm, inset 5 µm. **(H)** DIV14 cultured mouse cortical neurons co-expressing PLCXD2-FLAG and GFP were immunostained for FLAG (red), GFP (gray) and PSD95 (cyan). Scale bar 10 µm, inset 5 µm. **(I)** L4 barrel cortex neuron demonstrating endogenously tagged PLCXD2 expression at P21 as visualized by HA immunohistochemistry (left). Higher magnification inset shows prominent PLCXD2 localization in dendritic spines (right). Blue hues in the calibration bar indicate low expression, yellow hues indicate high expression. Scale bar 10 µm, inset 5 µm.

Binary MAPPIT experiments confirmed a specific interaction of PLCXD2 with the cytoplasmic domain of GPR158 **(Figure 1B)**. Single-molecule fluorescent in situ hybridization (smFISH) demonstrated that *Gpr158* and *Plcxd2* transcripts are co-expressed in excitatory L4 neurons, identified by *Rorb* expression **(Figure S1C)**. Both transcripts are enriched in L4 of the barrel field as early as postnatal day 4 (P4) and remain co-expressed up to P14 when L4 circuitry is fully established **(Figure S1D)**. Interestingly, in the developing hippocampus, *Plcxd2* and *Gpr158* are similarly co-expressed in CA3 neurons **(Figure S1E)**.

To validate the interaction, we reciprocally co-immunoprecipitated FLAG-tagged PLCXD2 together with HA-tagged GPR158 from HEK239T cell lysates **(Figure 1C)**. Live immunolabeling of the HA epitope in transfected HEK239T cells to specifically detect the surface pool of GPR158 showed clear colocalization with PLCXD2 at the PM, especially evident in orthogonal planes **(Figure 1D)**. Finally, we used split fluorescence-activating and absorption shifting tag (FAST), a reversible fluorescence complementation system, to visualize the interaction of GPR158 with PLCXD2 in live cells **(Figure 1E)**. SplitFAST relies on the binding of hydroxybenzylidene rhodanine (HBR) analogs that are not fluorescent in solution, but strongly fluoresce when immobilized in the binding cavity of FAST.^38^ GPR158-CFAST and PLCXD2-NFAST were co-expressed in HEK239T cells and fluorescence was live monitored with or without the addition of HMBR, an HBR analog. In the presence of HMBR, FAST fluorescence was almost exclusively present at the periphery of the cell, confirming that the GPR158-PLCXD2 complex resides at the PM **(Figure 1F)**. To determine whether splitFAST reliably reports protein-protein interactions, we co-expressed GPR158 and its only known intracellular binding partner RGS7 with or without Gβ5. In the absence of Gβ5, RGS7 is not able to bind to GPR158.^31^ Only when GPR158- CFAST and RGS7-NFAST were co-expressed together with Gβ5, FAST fluorescence was visible, thus excluding possible spontaneous self-assembly of the FAST fragments **(Figure S2A)**. When we co-expressed GPR158-CFAST and PLCXD2-NFAST in cultured cortical neurons, FAST fluorescence was observed in a punctate pattern along dendrites and in dendritic spines **(Figure 1G)**.

To assess the subcellular distribution of PLCXD2, we transfected PLCXD2-FLAG in cultured cortical neurons, as both commercially available and two custom-made PLCXD2 antibodies were non-specific. This demonstrated a prominent punctate localization of PLCXD2 adjacent to the excitatory postsynaptic density protein PSD95 in some, but not all dendritic spines **(Figure 1H)**. To circumvent possible overexpression-related artefacts, we next endogenously tagged PLCXD2 with spaghetti monster HA (smHA) using a CRISPR/Cas9-mediated homology-independent targeted integration strategy.^39^ Following adeno-associated virus (AAV)- based delivery of the knock-in construct in cultured cortical neurons from *H11^Cas9^* mice, we noted a similar punctate localization of PLCXD2 in some, but not all, dendritic spines and along the dendritic PM. PLCXD2 was exclusively detected in the somatodendritic compartment **(Figure S2B)**. To examine the subcellular distribution of PLCXD2 *in vivo*, we injected AAVs harboring the knock-in cassette in the barrel cortex of *H11^Cas9^* mice. Endogenously tagged PLCXD2 showed a punctate distribution in dendritic spines of L4 cortical neurons **(Figure 1I, Figure S2C)**. Using a KO validated antibody^26^, we confirmed postsynaptic localization of GPR158 in L4 neurons, at thalamocortical (marked by VGluT2) and intracortical (marked by VGluT1) synapses **(Figure S2D)**. Together, these results identify a novel GPR158-PLCXD2 complex that localizes to the postsynaptic compartment.

### GPR158 prevents PLCXD2-induced ER-PM contact site remodeling

PLCXD2 is an atypical PLC that was recently identified as one of three members from the PLCXD family. It has no known function in the brain. In humans, three PLCXD2 isoforms exist, of which hPLCXD2.1 is reported to localize to the nucleus. Overexpression of hPLCXD2.1 in heterologous cells increases the turnover of radiolabeled inositol phosphate, indicating enhanced basal PLC activity.^40^ In invertebrates, the *Drosophila* orthologue dPLCXD was found to control PIP2 levels on endolysosomal membranes.^41^ To obtain clues about the function of mouse PLCXD2, we first investigated its subcellular localization in HEK239T cells together with a panel of organelle markers. In contrast to hPLCXD2.1 and dPLCXD, mPLCXD2 showed profound PM labeling with no evidence of nuclear or endosomal accumulation **(Figures S3A and S3B)**. Total internal reflection fluorescence structured illumination microscopy (TIRF-SIM) imaging revealed the presence of PLCXD2 puncta interspersed between cortical ER-labeled structures near the PM **(Figure 2A)**. This prompted us to explore whether PLCXD2 might reside in close proximity to ER-PM contact sites. TIRF-SIM imaging of PLCXD2 together with MAPPER, a genetically encoded ER-PM contact site marker^42^, demonstrated prominent localization of PLCXD2 near these contact sites **(Figure 2B)**. Nearest-neighbor analysis of reconstructed puncta showed a close association between MAPPER and PLCXD2 that differed significantly from a dataset where original image channels were unpaired and randomly assigned to each other **(Figures 2C and 2D)**. A close spatial association was also found between PLCXD2 and extended synaptotagmin 2 (E-SYT2), an ER-resident tether that constitutively localizes to ER-PM contact sites^43^ **(Figure S3C)**.

**Figure 2.**
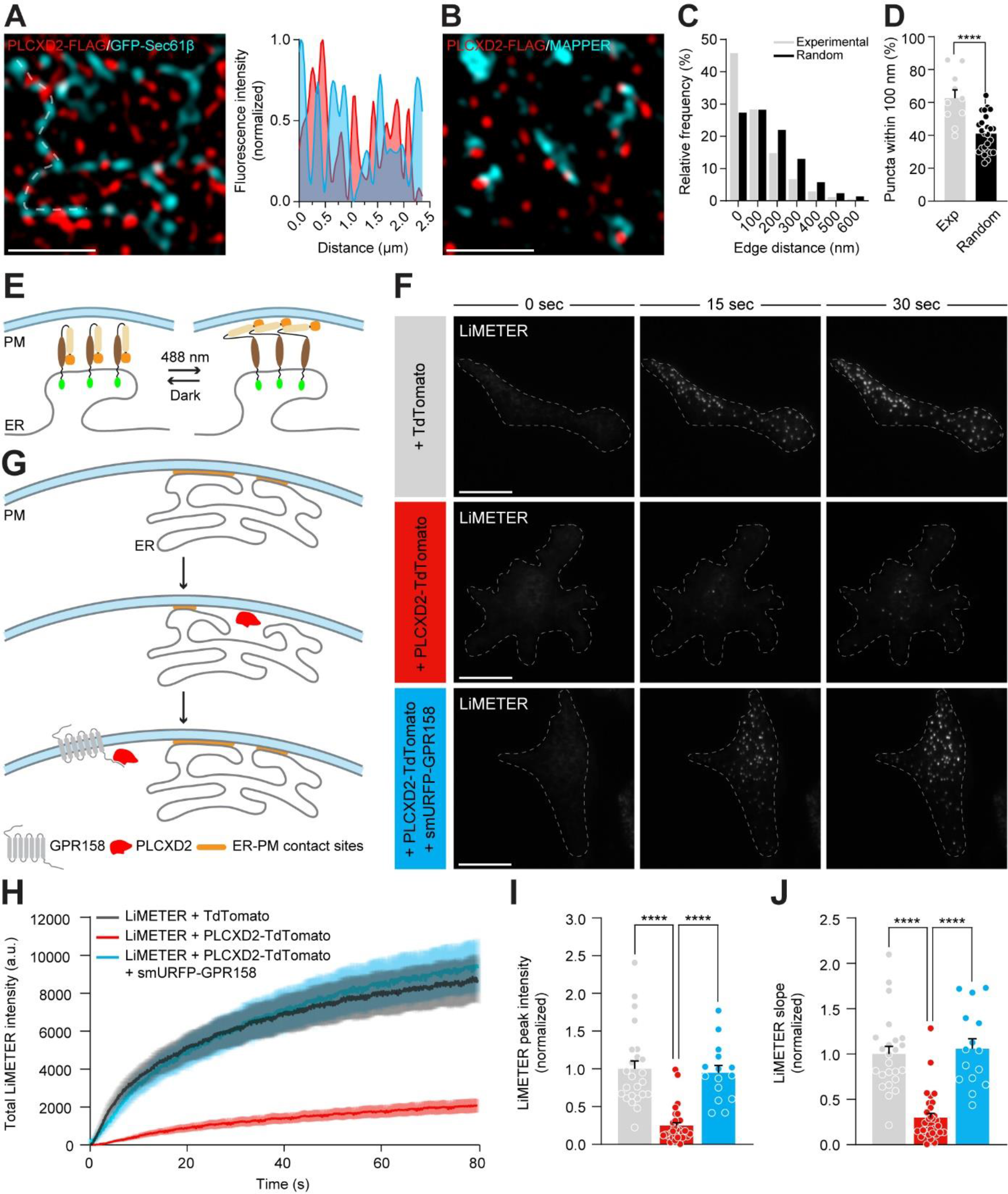
GPR158 prevents PLCXD2-induced ER-PM contact site remodeling (see also Supplementary figures S3 and S4). **(A)** TIRF-SIM images of HEK239T cells co-expressing PLCXD2-FLAG and GFP- Sec61β immunostained for FLAG (red) and GFP (cyan) (left). Fluorescence intensity profile line scans show PLCXD2 interspersed between cortical ER-labeled structures. Scale bar 1 µm. **(B)** TIRF-SIM images of HEK239T cells co-expressing PLCXD2-FLAG and MAPPER immunostained for FLAG (red) and GFP (cyan). Scale bar 1 µm. **(C)** Frequency distribution of PLCXD2-to-MAPPER puncta edge distances obtained with nearest- neighbor analysis. Random distances were calculated using unpaired experimental data (see Methods). **(D)** Quantification of experimental and random PLCXD2-to-MAPPER puncta edge distances below 100 nm (*n* = 10 cells). \*\*\**P*<0.001, two-tailed unpaired t-test. **(E)** Schematic representation of LiMETER, a photo-activatable ER-PM contact site marker. LiMETER contains a luminal GFP reporter (green) and the transmembrane domain of STIM1 (black). The cytoplasmic region consists of a light-sensitive LOV2 (brown)-Jα (beige) domain and the PM-targeting polybasic domain of STIM1 (orange). Blue light stimulation promotes unwinding of the Jα helix and exposes the polybasic domain of STIM1 to enable interaction with the PM. **(F)** Representative TIRF images of HEK239T cells co-expressing LiMETER either with TdTomato (control), with PLCXD2-TdTomato or with PLCXD2-TdTomato and smURFP-GPR158 at the indicated timepoints. Scale bar 10 µm. **(G)** Schematic representation of the interplay between GPR158 and PLCXD2 during ER-PM contact site formation. GPR158-mediated PLCXD2 inhibition prevents continuous PIP2 depletion and allows for ER-PM contact site formation to proceed. **(H)** Time course of LiMETER accumulation following blue light stimulation in HEK239T cells co-expressing LiMETER and TdTomato (black trace), co-expressing LiMETER and PLCXD2-TdTomato (red trace) or co-expressing LiMETER, PLCXD2-TdTomato and smURFP-GPR158 (cyan). **(I and J)** Quantification of the peak LiMETER intensity **(I)** and the rate of LiMETER accumulation **(J)**. LiMETER + TdTomato (*n* = 3 experiments, 24 cells), LiMETER + PLCXD2-TdTomato (*n* = 3 experiments, 31 cells), LiMETER + PLCXD2-TdTomato + smURFP-GPR158 (*n* = 3, experiments 15 cells). \*\*\*\**P*<0.0001, Kruskal-Wallis test, Dunn’s multiple comparisons.

Compared to all other PLCs, PLCXD2 only contains an X-domain that is part of the catalytic X-Y barrel, and is devoid of any other regulatory domains.^40^ Importantly, two histidine residues critical for the enzymatic activity of PLCs are evolutionarily conserved **(Figure S4A)**, suggesting that PLCXD2 might retain enzymatic activity. Indeed, expression of PLCXD2, but not of a catalytic dead mutant (PLCXD2 CD) in which these histidine residues were replaced with leucines, led to a reduction in PM PIP2 levels indicated by a translocation of the genetically encoded PIP2 probe PH- PLCδ-GFP from the PM to the cytosol **(Figures S4B and S4C)**. Co-expression of PLCXD2 with GPR158 prevented translocation of PH-PLCδ-GFP to the cytosol, indicating that binding to GPR158 inhibits the enzymatic activity of PLCXD2 **(Figures S4D and S4E)**.

The stable formation of ER-PM contact sites heavily relies on PIP2 levels in the PM. Several ER-PM tethers as well as the ER Ca^2+^ sensor stromal interaction molecule 1 (STIM1), which initiates the formation of ER-PM contact sites, critically depend on PIP2 for their interaction with the PM.^44–49^ Using a light-inducible variant of MAPPER, termed LiMETER^50^ **(Figure 2E)**, we monitored the accumulation of ER-PM contact sites in HEK239T cells co-expressing PLCXD2 or both PLCXD2 and GPR158 using TIRF microscopy. Following blue light stimulation, LiMETER rapidly translocated to the PM in the form of discrete puncta **(Figures 2F-J, Movie S1)**. Compared to HEK239T cells expressing LiMETER alone, introducing PLCXD2 led to a drastic reduction in both the rate and extent of LiMETER accumulation at the PM **(Figures 2F-J, Movie S1)**. Co-expression of PLCXD2 and GPR158 fully rescued LiMETER accumulation, further confirming that GPR158 inhibits PLCXD2 function **(Figures 2F-2J, Movie S1)**. Live TIRF imaging of MAPPER to examine to stability of individual ER-PM junctions indicated a strong increase in the lateral mobility of MAPPER puncta when co-expressed with PLCXD2 **(Movies S2-S5)**.

Given that PLCXD2 impedes the formation of ER-PM contact sites, we wondered whether PLCXD2 also impairs store operated calcium entry (SOCE). SOCE is initiated by depletion of Ca^2+^ from the ER and triggers a series of events that culminate in the formation of ER-PM contact sites.^51^ Here, STIM proteins in the ER and Ca^2+^-selective ORAI channels in the PM interact to allow Ca^2+^ influx across the PM.^52,53^ HEK239T cells were treated with the sarcolemma ER Ca^2+^ ATPase (SERCA) inhibitor cyclopiazonic acid (CPA) in a Ca^2+^-free solution to deplete ER Ca^2+^ stores. The SOCE response was then assessed by analyzing cytosolic Ca^2+^ influx following re-addition of extracellular Ca^2+^. The rate and extent of the SOCE response were strongly impaired in PLCXD2 expressing cells compared to surrounding non-transfected cells **(Figures S4F, S4H and S4I)**. In agreement with GPR158 inhibiting PLCXD2, we observed no difference in the SOCE response between PLCXD2 and GPR158 co-expressing cells and surrounding non-transfected cells **(Figures S4G-S4I)**. These findings demonstrate that PLCXD2 acts as a constitutively active PLC residing in close proximity to ER-PM contact sites. Binding of PLCXD2 to GPR158 prevents continuous PIP2 depletion, allowing for the formation of stable ER-PM contact sites and for SOCE to proceed.

### PLCXD2 hinders SA formation

Our data derived from heterologous cells show the importance of control over PIP2 levels for the proper formation and function of ER-PM contact sites. In the postsynaptic compartment, a modification of the local membrane lipid microenvironment could have several consequences, including altering the behavior of ER-PM tethering proteins. This may hamper the formation of SA-related ER-PM contacts, which are readily observed in dendritic spines at the electron microscopy level^4,5,22–24^, and hinder SA stabilization. On the other hand, PIP2 is a well-known F-actin regulator promoting actin polymerization.^54^ Recent work alluded to the critical importance of an intact spinous F-actin network, providing the required tracks for myoVa to stably anchor an sER tubule and facilitate SA formation.^18,19^ Thus, we reasoned that elevated levels of PLCXD2 in dendritic spines would create an unfavorable environment for SA formation.

We first confirmed that PLCXD2 locally depletes PIP2 in dendritic spines. PLCXD2 demarcated regions in dendritic spines of cultured hippocampal neurons that were devoid of PH-PLCδ-GFP fluorescence **(Figure S5)**. In agreement with PIP2 regulating actin polymerization, F-actin fluorescence intensity was reduced in PLCXD2- containing spines compared to spines lacking PLCXD2 **(Figures 3A-3C)**. Immunolabeling for endogenous synaptopodin (SYNPO), the main structural component of the SA and a well-established marker for this organelle^6^, revealed that PLCXD2 and SYNPO are rarely present in the same spine, demonstrating an almost mutually exclusive localization pattern **(Figures 3D and 3E)**. Furthermore, in the rare occasions where PLCXD2 and SYNPO were found together in the same spine, SYNPO fluorescence intensity was strongly diminished compared to spines lacking PLCXD2 **(Figure 3F)**. Thus, PLCXD2 creates an unfavorable environment that hinders SA formation, presumably via limiting the availability of PIP2 and interfering with the accumulation of spinous F-actin.

**Figure 3.**
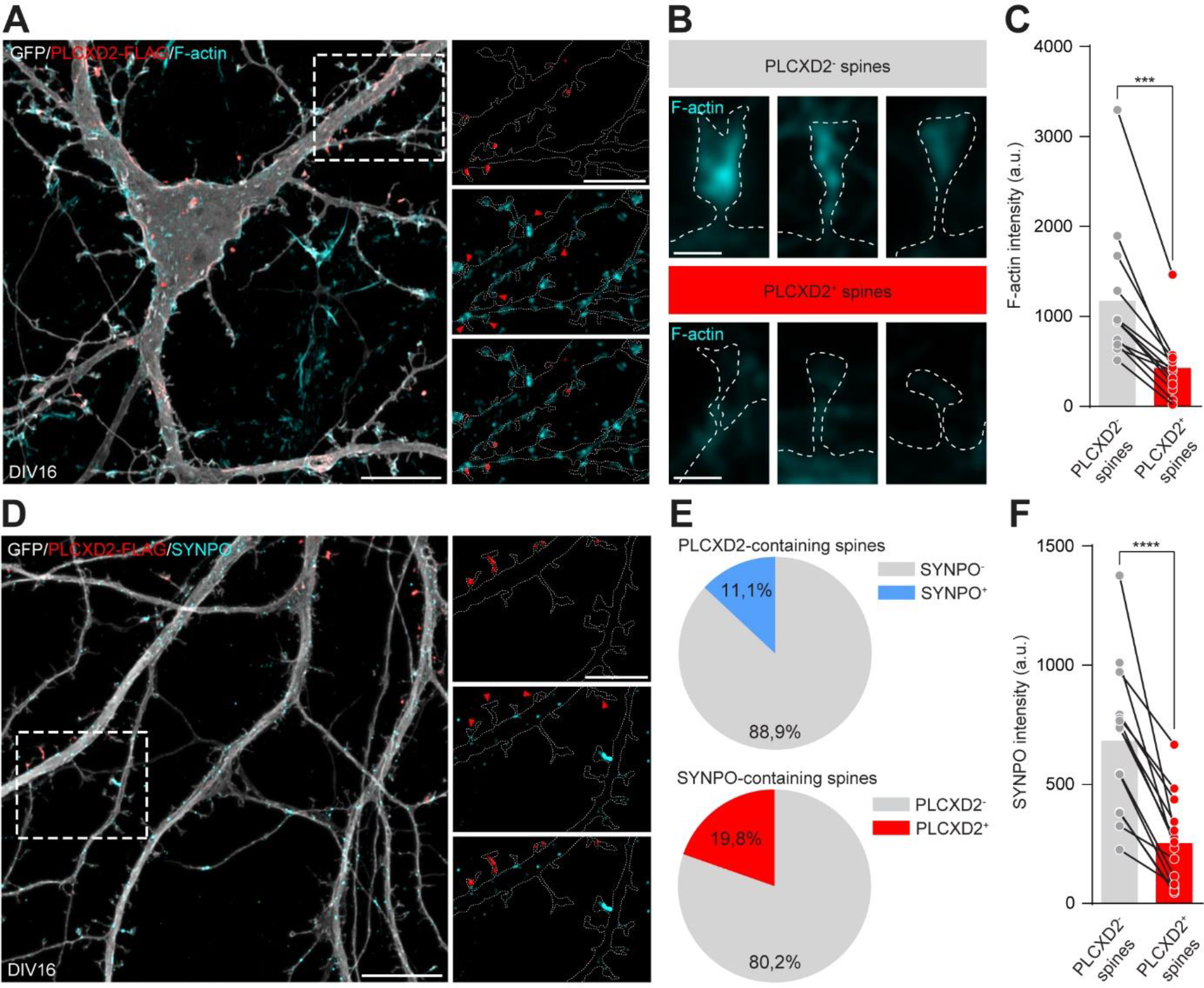
PLCXD2 hinders SA formation (see also Supplementary figure S5). **(A)** DIV16 mouse cultured hippocampal neuron co-expressing PLCXD2-FLAG and GFP immunostained for FLAG (red), GFP (gray) and F-actin (cyan). Scale bar 10 µm, inset 5 µm. **(B)** Representative images of dendritic spines from cultured hippocampal neurons lacking or containing PLCXD2-FLAG, immunostained for F-actin (cyan). Scale bar 1 µm. **(C)** Quantification of F-actin fluorescence intensity in spines. Each pair of dot plots represents the average F-actin fluorescence intensity in PLCXD2- and PLCXD2+ spines from the same neuron (*n* = 3 cultures, 13 neurons). \*\*\**P*<0.001, Wilcoxon matched-pairs signed rank test. **(D)** DIV16 mouse cultured hippocampal neuron co-expressing PLCXD2-FLAG and GFP were immunostained for FLAG (red), GFP (gray) and SYNPO (cyan). Scale bar 10 µm, inset 5 µm. **(E)** Percentage of PLCXD2+ spines containing SYNPO and vice versa. **(F)** Quantification of SYNPO fluorescence intensity. Each pair of dot plots represents the average SYNPO fluorescence intensity in PLCXD2- and PLCXD2+ spines from the same neuron (*n* = 3 cultures, 14 neurons). \*\*\*\**P*<0.0001, two-tailed paired *t*-test.

### A GPR158-PLCXD2 interaction regulates SA content and maturation of dendritic spines

Several studies have demonstrated the importance of the SA for the proper maturation of dendritic spines.^13,16,55^ Given the striking co-expression of *Gpr158* and *Plcxd2* **(Figures S1D and S1E)** and their interaction in the postsynaptic compartment **(Figure 1G)**, we speculated that loss of *Gpr158*, and thus unrestricted PLCXD2 activity would impair dendritic spine maturation. We generated *Gpr158* floxed mice **(Figures S6A and S6B)** and conditionally deleted *Gpr158* from excitatory L4 cortical neurons using the *Rorb-Cre* driver line.^56^ L4 cKO (*Gpr158^fl/fl^:Rorb-Cre*) and WT (*Gpr158^+/+^:Rorb-Cre*) littermate control mice were injected with a mixture of AAV-TRE-DIO-FLPo and AAV-TRE-fDIO-GFP-IRES-tTA in the barrel cortex at P0 to sparsely, but brightly label individual neurons^57^ **(Figure 4A)**. Dendritic spine morphology and density were analyzed at P14. Compared to WT neurons, dendritic spines from *Gpr158* L4 cKO neurons were shifted towards a less mature phenotype. Loss of GPR158 led to a decrease in the fraction of mature mushroom-type spines with a concomitant increase in immature filopodia-type protrusions **(Figures 4B and 4C)**. Dendritic spine density was not affected in the absence of GPR158 **(Figure 4D)**.

**Figure 4.**
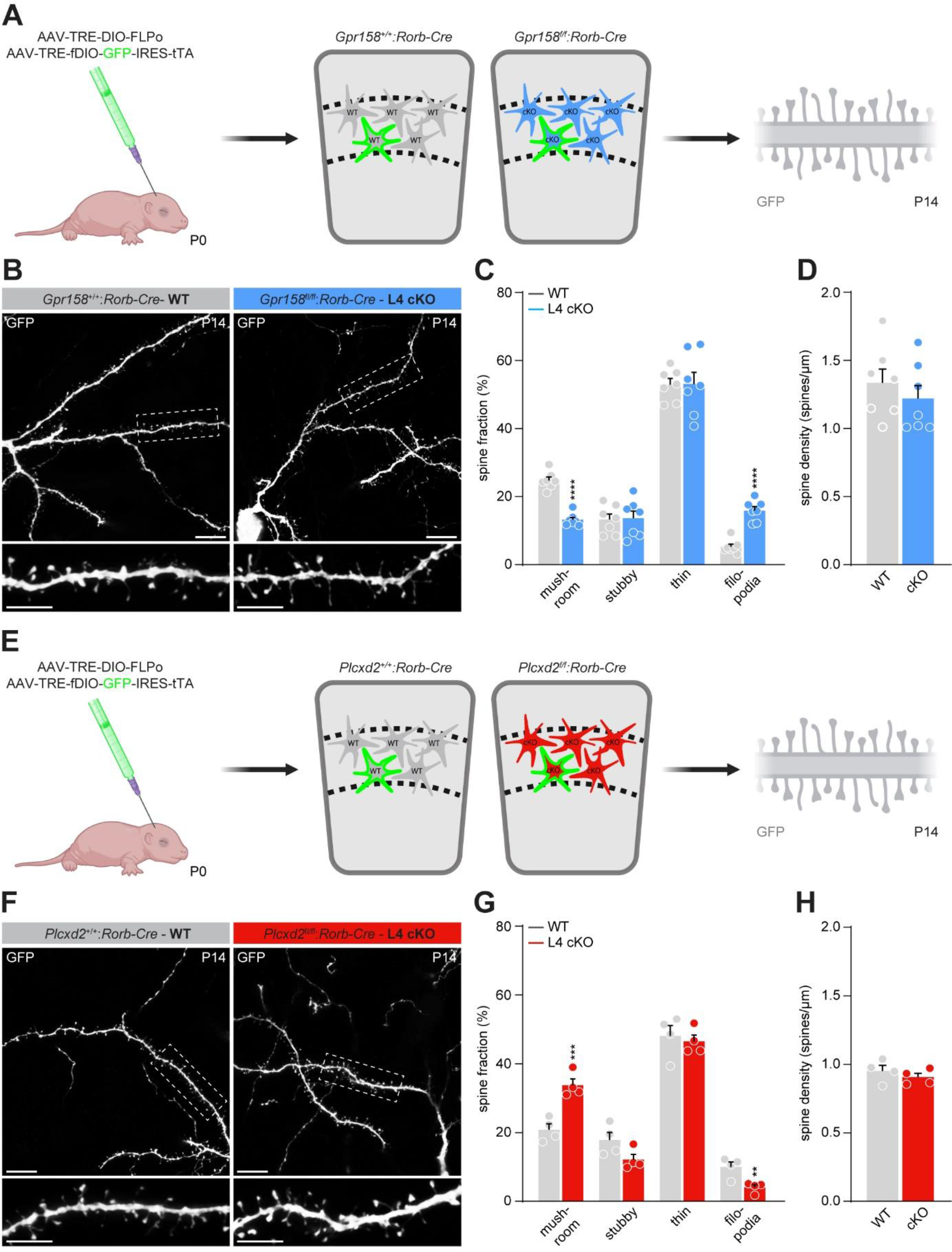
GPR158 and PLCXD2 oppositely regulate dendritic spine maturation (see also Supplementary figure S6). **(A)** Experimental strategy to achieve sparse, but bright labeling of *Gpr158* WT (*Gpr158^+/+^:Rorb-Cre*) and *Gpr158* L4 cKO (*Gpr158^fl/fl^:Rorb-Cre*) neurons. In Cre-positive neurons, successive tTA-TRE cycles facilitate expression of FLPo and GFP in the scarce population of transduced neurons. **(B)** Representative images of *Gpr158* WT and L4 cKO neurons in the barrel cortex, immunostained for GFP at P14. Note the appearance of numerous immature filopodia in *Gpr158* cKO neurons. Scale bar 10 µm, inset 5 µm. **(C)** Quantification of dendritic spine morphology in *Gpr158* WT (*N* = 7 mice, *n* = 19 neurons) and L4 cKO neurons (*N* = 7 mice, *n* = 25 neurons). \*\*\*\**P*<0.0001, nested two- tailed unpaired *t*-test. **(D)** Quantification of dendritic spine density in *Gpr158* WT (*N* = 7 mice, *n* = 19 neurons) and L4 cKO neurons (*N* = 7 mice, *n* = 25 neurons). **(E)** Experimental strategy to achieve sparse, but bright labeling of *Plcxd2* WT (*Plcxd2^+/+^:Rorb-Cre*) and Plcxd2 L4 cKO (*Plcxd2^fl/fl^:Rorb-Cre*) neurons. In Cre-positive neurons, successive tTA-TRE cycles facilitate expression of FLPo and GFP in the scarce population of transduced neurons. **(F)** Representative images of *Plcxd2* WT and L4 cKO neurons in the barrel cortex, immunostained for GFP at P14. **(G)** Quantification of dendritic spine morphology in *Plcxd2* WT (*N* = 4 mice, *n* = 10 neurons) and L4 cKO neurons (*N* = 4 mice, *n* = 9 neurons). \*\**P*<0.01, \*\*\**P*<0.001, nested two-tailed unpaired *t*-test. Scale bar 10 µm, inset 5 µm. **(H)** Quantification of dendritic spine density in *Plcxd2* WT (*N* = 4 mice, *n* = 10 neurons) and L4 cKO neurons (*N* = 4 mice, *n* = 9 neurons).

We next investigated how loss of PLCXD2 would influence the maturation of the postsynaptic compartment. Since PLCXD2 hinders SA formation **(Figure 3)**, we predicted that removing PLCXD2 would promote dendritic spine maturation. We generated *Plcxd2* floxed mice **(Figures S6C and S6D)** and used the same strategy to conditionally delete *Plcxd2* in L4 cortical neurons **(Figure 4E)**. In contrast to neurons lacking GPR158, dendritic spines of neurons lacking PLCXD2 were indeed shifted towards a more mature phenotype **(Figure 4F)**. Compared to WT (*Plcxd2^+/+^:Rorb-Cre*) mice, neurons from *Plcxd2* L4 cKO (*Plcxd2^fl/fl^:Rorb-Cre*) mice exhibited fewer immature filopodia-type protrusions while mature mushroom-type spines were more numerous **(Figure 4G)**. Dendritic spine density was similar between *Plcxd2* WT and L4 cKO neurons **(Figure 4H)**. Together, these *in vivo* observations are consistent with our *in vitro* data showing GPR158 inhibitory control over PLCXD2 function. In the absence of GPR158, unchecked PLCXD2 activity attenuates dendritic spine maturation. Conversely, removing PLCXD2 promotes dendritic spine maturation, possibly by alleviating a break on SA incorporation.

To exclude any non-cell autonomous phenotypes that may be associated with the removal of GPR158 or PLCXD2 across the entirety of L4, we next opted for a system in which gene expression could be manipulated in a sparse manner. A mixture of AAV-TRE-Cre and AAV-SYN-DIO-GFP-IRES-tTA was injected in the barrel cortex of WT control mice (*Gpr158^+/+^:Plcxd2^+/+^*), *Gpr158* cKO mice (*Gpr158^f/f^:Plcxd2^+/+^*) and *Gpr158:Plcxd2* double cKO mice (*Gpr158^f/f^:Plcxd2^f/f^*) at P0. Using this strategy, only a small population of neurons will express high levels of Cre and GFP, allowing simultaneous gene cKO and cell labeling^58,59^ **(Figure 5A)**. Following two weeks of expression, sections were immunostained for GFP and endogenous SYNPO to analyze the morphology and density of dendritic spines, and assess their SA occupancy **(Figure S7)**. Removal of GPR158 in a sparse population of L4 cortical neurons recapitulated the shift towards an immature spine morphology **(Figures 5B, 5C and 5E)** we observed following L4-wide deletion of *Gpr158* and led to a strong reduction in the number of SA-bearing spines **(Figures 5B, 5C and 5F)**. Importantly, this loss of SA-bearing spines was not simply due to a reduction in the number of mature spines, the type of spines that the SA is tightly associated with.^5–7^ We found that 40-60% of mushroom-type spines contained a spine apparatus in WT neurons, while this fraction was halved in the remaining mushroom-type spines of *Gpr158* cKO neurons **(Figures 5B, 5C and 5G)**.

**Figure 5.**
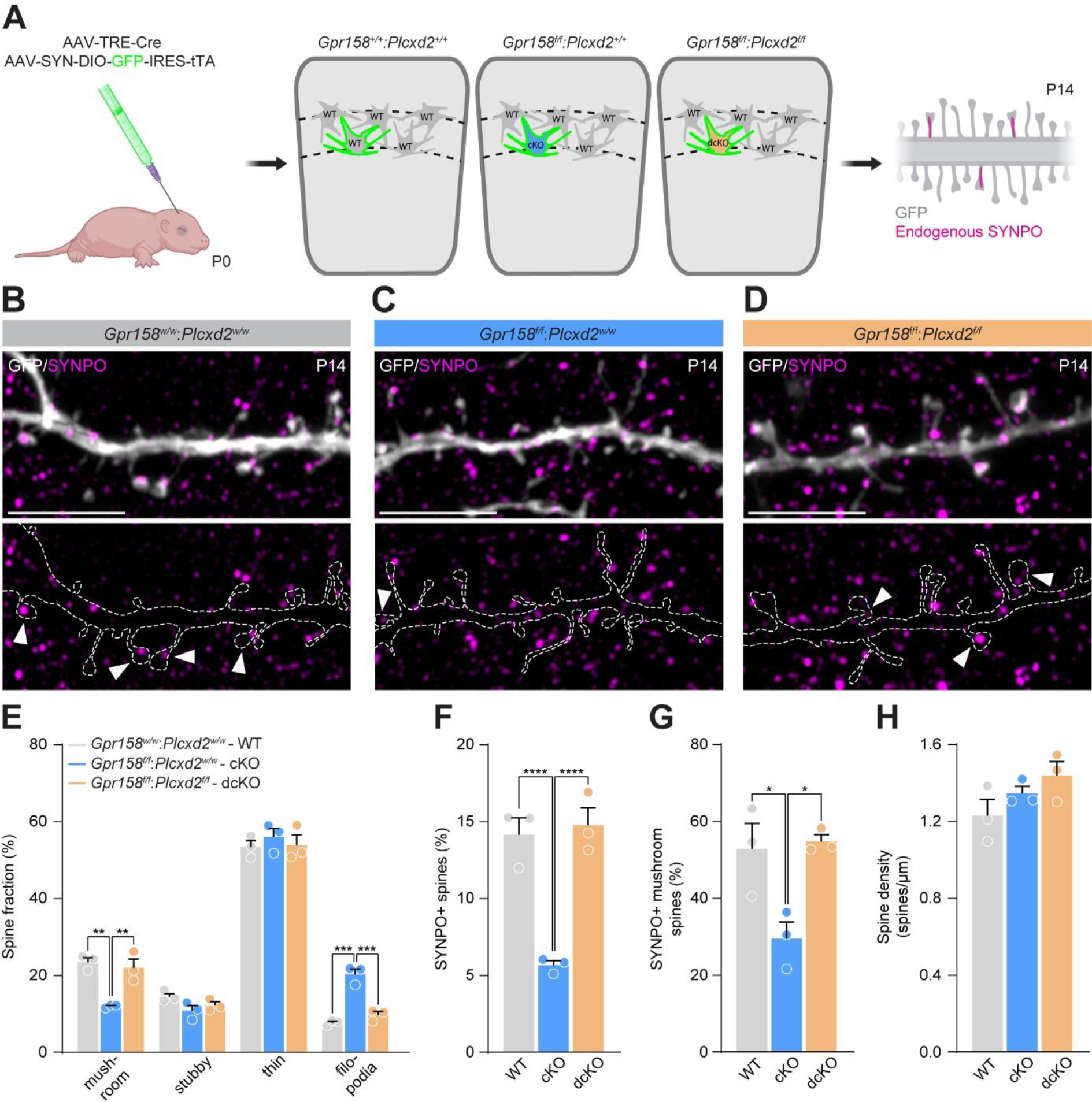
A GPR158-PLCXD2 complex regulates SA content and maturation of dendritic spines (see also Supplementary figure S7). **(A)** Experimental strategy to achieve simultaneous sparse gene cKO and cell labeling. Leakage of the TRE promoter drives weak expression of Cre and, subsequently, tTA in a very small population of neurons that carry both vectors. Next, tTA enhances Cre expression by binding to TRE. This positive feedback of successive tTA-TRE cycles facilitates expression of Cre and GFP in the scarce population of transduced neurons. **(B-D)** Representative images of dendritic segments from WT **(B)**, *Gpr158* cKO **(C)** and *Gpr158*:*Plcxd2* dcKO **(D)** neurons in L4 of the barrel cortex, immunostained for GFP and SYNPO at P14. Scale bar 5 µm. (**E)** Quantification of dendritic spine morphology in WT (N = 3 mice, n = 9 neurons), *Gpr158* cKO (*N* = 3 mice, *n* = 9 neurons) and *Gpr158*:*Plcxd2* dcKO neurons (*N* = 3 mice, *n* = 9 neurons). \*\**P*<0.001, \*\*\**P*<0.001, nested 1-way ANOVA, Tukey’s multiple comparisons. **(F)** Quantification of SYNPO-containing spines in WT (N = 3 mice, n = 9 neurons), *Gpr158* cKO (*N* = 3 mice, *n* = 9 neurons) and *Gpr158*:*Plcxd2* dcKO neurons (*N* = 3 mice, *n* = 9 neurons). \*\*\*\**P*<0.0001, nested 1-way ANOVA, Tukey’s multiple comparisons. **(G)** Quantification of SYNPO-containing mushroom-type spines in WT (*N* = 3 mice, *n* = 9 neurons), *Gpr158* cKO (*N* = 3 mice, *n* = 9 neurons) and *Gpr158*:*Plcxd2* dcKO neurons (*N* = 3 mice, *n* = 9 neurons). \**P*<0.05, nested 1-way ANOVA, Tukey’s multiple comparisons. **(H)** Quantification of dendritic spine density in WT (*N* = 3 mice, *n* = 9 neurons), *Gpr158* cKO (*N* = 3 mice, *n* = 9 neurons) and *Gpr158*:*Plcxd2* dcKO neurons (*N* = 3 mice, *n* = 9 neurons).

To determine whether the shift towards immature dendritic spine morphology and a reduction of SA-bearing spines in the absence of GPR158 depends on PLCXD2, we next assessed spine morphology and SA occupancy in *Gpr158:Plcxd2* double cKO mice. The loss of mature mushroom-type spines and an increase in immature filopodia-type protrusions in *Gpr158* cKO neurons was rescued in *Gpr158:Plcxd2* double cKO neurons **(Figures 5B, 5D and 5E)**. Moreover, removal of *Plcxd2* in *Gpr158* cKO neurons rescued SA content of dendritic spines **(Figures 5B, 5D, 5F and 5G)**, demonstrating that in the absence of GPR158, unrestrained PLCXD2 activity prevents postsynaptic SA incorporation. Spine density was not altered in either condition **(Figure 5H)**. Taken together, these findings indicate that GPR158 inhibitory control over PLCXD2 cell-autonomously regulates SA content and the maturation of dendritic spines.

### Extracellular HSPG binding modulates the GPR158-PLCXD2 complex

The extracellular domain of GPR158 interacts with the HS-moiety of HSPGs, including glypicans (GPCs)^29^ that may act as presynaptic GPR158 binding partners.^26^ To determine whether the GPR158-PLCXD2 interaction might be modulated by binding of HSPGs to GPR158, we returned to the splitFAST system. HEK239T express HSPGs.^60^ We therefore reasoned that GPR158 expressed in these cells might interact with HSPGs in *cis* and that removal of HS using heparinases would abolish these interactions. HEK239T cells co-expressing GPR158-CFAST and PLCXD2-NFAST were treated with heparinase I/III for 2 hours to remove HS from the cell surface. We verified that heparinase I/III treatment did not alter the surface expression of GPR158 **(Figure S8A)** and efficiently removed HS from the cell surface **(Figure S8B)**. Compared to vehicle-treated cells, heparinase I/III-treated cells displayed a robust decrease in FAST fluorescence **(Figures 6A-C)**, indicating disassembly of the GPR158-PLCXD2 complex following removal of cell-surface HS. Subsequent application of recombinant GPC1 to the surface of heparinase I/III-treated cells augmented FAST fluorescence again **(Figures 6A-C)**. Together, these results indicate that cell-surface HPSGs modulate the GPR158-PLCXD2 interaction: removal of cell-surface HS disrupts the interaction, whereas the addition of an HSPG promotes it.

**Figure 6.**
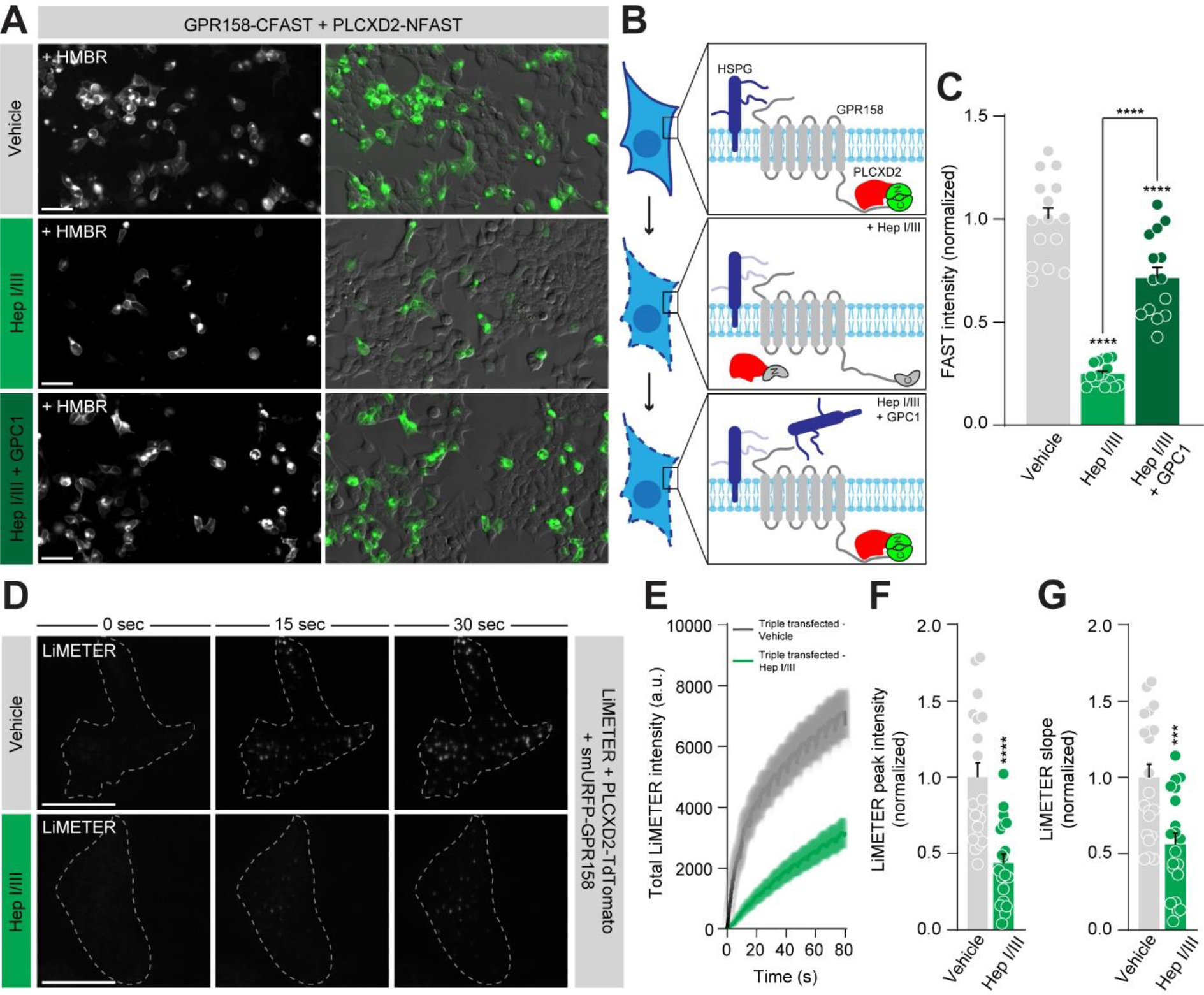
Extracellular HSPG binding modulates the GPR158-PLCXD2 complex (see also Supplementary figure S8). **(A)** HEK239T cells co-expressing GPR158-CFAST and PLCXD2-NFAST were visualized live in the presence of HMBR. Cells were either treated with vehicle or heparinase I/III. Recombinant GPC1 was added following 2 hours of heparinase I/III treatment. Scale bar 50 µm. **(B)** Schematic representation summarizing how HSPG binding may modulate the GPR158-PLCXD2 interaction. Disrupting extracellular HSPG-GPR158 binding by removing the HS-moieties of HSPGs using heparinase I/III disassembles the GPR158- PLCXD2 complex. Subsequent application of recombinant GPC1 promotes the GPR158-PLCXD2 interaction. **(C)** Quantification of FAST fluorescence intensity. Vehicle (*n* = 4 experiments, 15 wells), heparinase I/III (*n* = 4 experiments, 16 wells), heparinase I/III + GPC1 (*n* = 4 experiments, 15 wells). \*\*\*\**P*<0.0001, 1-way ANOVA, Bonferroni’s multiple comparisons. **(D)** Representative TIRF images of HEK239T cells co-expressing LiMETER, PLCXD2- TdTomato and smURFP-GPR158. Cells were treated with vehicle or heparinase I/III for 2 hours. Scale bar 10 µm. **(E)** Time course of LiMETER accumulation following blue light stimulation in HEK239T cells co-expressing LiMETER, PLCXD2-TdTomato and smURFP-GPR158 treated with vehicle (black trace) or heparinase I/III (green trace). **(F and G)** Quantification of peak LiMETER intensity **(F)** and rate of LiMETER accumulation **(G)** in cells treated with vehicle (*n* = 3 experiments, 21 cells) or heparinase I/III (*n* = 3 experiments, 20 cells). \*\*\*\**P*<0.0001, Mann-Whitney test, \*\*\**P*<0.001, two-tailed unpaired t-test.

Lastly, we considered that dissociation of the GPR158-PLCXD2 interaction by removing cell-surface HS, and thus eliminating GPR158 inhibitory control over PLCXD2, would limit the formation of ER-PM contact sites. We therefore co-expressed GPR158 and PLCXD2 in HEK239T cells and evaluated LiMETER accumulation following removal of cell-surface HS. Indeed, LiMETER translocation to the PM assessed by TIRF microscopy was strongly reduced following heparinase I/III treatment compared to vehicle-treated cells **(Figures 6D-6G)**, indicating an impediment in the formation of ER-PM contact sites. Thus, HSPG binding modulates the GPR158-PLCXD2 association and downstream function.

## Discussion

The molecular mechanisms that ensure the specific localization of the SA at a subset of dendritic spines are largely unknown. Here, we show that the postsynaptic cell-surface receptor GPR158 cell-autonomously signals via PLCXD2 to control SA abundance and the maturation of dendritic spines **(Figure S9)**.

We identify PLCXD2, an unconventional and poorly characterized PLC with no known function in the brain, as a novel intracellular GPR158 binding partner. PLCXD2 lacks all known regulatory domains that are found in classical PLCs. Despite lacking a pleckstrin homology (PH)-domain or C2 domain that can mediate membrane binding^61,62^, our findings demonstrate prominent PM localization for PLCXD2. Posttranslational modifications such as S-palmitoylation play an important role in targeting proteins to the PM, including numerous synaptic proteins.^63^ Indeed, pCysMod, a deep learning-based tool that identifies cysteine modifications^64^ (http://pcysmod.omicsbio.info/index.php), predicts several high-confident cysteine residues in PLCXD2 that are potentially S-palmitoylated and may provide an explanation for its membrane targeting. PLCXD2 harbors a single X-domain, part of the catalytic X-Y barrel.^40^ Importantly, an X-Y linker, absent in PLCXD2, maintains classical PLCs in an auto-inhibited state.^61,62^ This suggests the existence of additional mechanisms to control its enzymatic activity. Accordingly, we find that PLCXD2 constitutively hydrolyses PIP2 and is inhibited by binding to GPR158. The X-Y linker in classical PLCs adopts a helical shape, blocking access to the enzyme’s active site.^65,66^ The cytoplasmic domain of GPR158 consists of several α-helices^31,32^, one of which might perform a similar role and mask the active site of PLCXD2.

This mode of PLCXD2 regulation differs substantially from RGS7-mediated signaling in which GPR158 acts as a membrane anchor that recruits and potentiates RGS7’s GTPase activity at the PM.^31^ Whereas loss of GPR158 in CA3 hippocampal neurons^26^ and L4 cortical neurons (this study) hinders dendritic spine maturation, loss of GPR158 in layer 2/3 mPFC neurons augments synaptic strength.^27^ Remarkably, *Gpr158* and *Plcxd2* are strongly co-expressed in CA3 hippocampal and L4 cortical neurons, but not in mPFC neurons. While *Gpr158* expression is enriched in L2/3 neurons of the mPFC^27^, *Plcxd2* expression is enriched in L5 mPFC neurons^67^, suggesting that GPR158 acts in a PLCXD2-independent manner in layer 2/3 mPFC neurons. It is thus tempting to speculate that the molecular context in which GPR158 operates dictates its role during synapse development. Interestingly, *Gpr158* and *Plcxd2* show a striking co-expression pattern in a select population of other neuron types, including mitral cells of the olfactory bulb and dopaminergic neurons of the pars compacta in the substantia nigra (Allen Brain Atlas), where they may operate in a similar manner to regulate postsynaptic development. Which regions in GPR158’s cytoplasmic domain are responsible for interacting with PLCXD2 and whether RGS7 and PLCXD2 might compete for binding to GPR158 remain to be addressed in future work.

Our data in heterologous cells demonstrate that PLCXD2 associates with ER-PM contact sites where sustained PIP2 depletion limits their formation and function. In neurons, PLCXD2 localizes to dendritic spines where it interferes with the accumulation of F-actin and hinders SA formation. *In vivo*, we show that sparse conditional deletion of *Gpr158* in L4 cortical neurons leads to a loss of SA-bearing dendritic spines. Removal of *Plcxd2* in *Gpr158* cKO neurons rescues SA abundance, supporting GPR158 inhibitory control over PLCXD2. We envision two non-mutually exclusive mechanisms via which PLCXD2 is involved in the regulation of SA localization. First, an intact spinous F-actin network provides the required tracks for MyoVa to stably anchor an sER tubule and facilitate SA formation.^18,19^ In addition, a recent proximity biotinylation-based proteomic screen identified numerous actin-based scaffolds that co-assemble together with SYNPO, further highlighting the functional association between the spinous actin cytoskeleton and the SA.^24^ Since PIP2 is a critical F-actin regulator that promotes actin polymerization^54^, continuous PIP2 depletion by PLCXD2 is likely to interfere with the formation of a SA. Second, most ER tethering proteins interact with PIP2 to maintain a close apposition between the ER and the PM, including E-Syts^43^, anoctamin 8 (ANO8)^68^, GRAM domain 2 (GRAMD2)^69^, junctophilins (JPH)^70^, oxysterol-binding protein (OSBP)-related proteins (ORPs)^71–73^ and TMEM24.^74,75^ Tethering proteins that have been identified in neurons predominantly localize to ER-PM contact sites that are present in the neuronal soma and proximal dendrites.^75–80^ Even though contacts between SA-related tubules and the PM are observed in nearly all SA-containing spines^4,5,22–24^, their molecular composition is mostly unknown. E-Syts have been detected at presynaptic compartments^81,82^, but their localization in dendritic spines is less clear. Nonetheless, knocking down E-Syt1 in cultured hippocampal neurons blocks AMPA receptor (AMPAR) surface accumulation following long-term potentiation (LTP).^83^ Moreover, reducing E-Syt1 levels *in vivo* prevents the increase in dendritic spine density that occurs when mice are housed in an enriched environment.^84^ These findings highlight the importance of E-Syt1 for the proper functioning of the postsynaptic compartment. Additionally, the tethering proteins TMEM24 and ORP2 were identified in the SYNPO proximity proteome.^24^ Together, these studies demonstrate that various tethering proteins identified in heterologous cells are present in dendritic spines, where they may perform a similar role and aid in the stabilization of the SA. Thus, impaired functioning of tethering proteins at SA-related ER-PM contact sites due to unrestrained PLCXD2 activity might contribute to ineffective SA formation.

We further demonstrate that GPR158-mediated inhibitory control over PLCXD2 activity regulates dendritic spine maturation. Several mechanisms have been proposed via which the SA may contribute to dendritic spine development. Growing evidence indicates that SOCE constitutes a key route of Ca^2+^ influx across the postsynaptic PM necessary to activate Ca^2+^-dependent signaling pathways that enable postsynaptic maturation and plasticity.^85^ In neurons, both STIM and ORAI proteins are detected at synapses and preferentially localize to SA-bearing spines.^55,86^ Loss of STIM2 or ORAI1 induces an impairment in dendritic spine maturation^87–90^ and attenuates LTP.^91^ In addition, STIM2 may directly interact with GluA1-containing AMPARs to control their plasticity-induced surface expression.^89,92^ Finally, the SA promotes synaptic AMPAR retention by limiting their surface diffusion.^11^

We show that binding of an HSPG to GPR158 modulates the GPR158-PLCXD2 association. Detailed analysis of cryo-electronmicroscopy structures from GPR158 in its apo state and in complex with RGS7/Gβ5 indicated allosteric coupling between GPR158’s extracellular and intracellular domain, and provided evidence for the existence of a potential small molecule ligand.^32,93^ Indeed, glycine was recently identified as an endogenous GPR158 ligand that directly binds to its extracellular Cache domain and reduces RGS7’s GTPase activity, possibly by disfavoring its orientation toward the PM or causing dissociation of RGS7/Gβ5 from GPR158.^30^ HSPGs interact with GPR158’s extracellular domain via their HS moiety^26,29^, which consists of repeated negatively charged polysaccharide units. The binding sites of HSPGs in the extracellular domain of GPR158 are not known but are unlikely to involve the small molecule-binding pocket in the Cache domain. Together with our finding that GPC1 promotes the GPR158-PLCXD2 interaction, these observations suggest that PLCXD2-mediated signaling is likely subject to a different mode of extracellular coupling, involving distinct structural features of GPR158. Whether glycine can regulate the GPR158-PLCXD2 interaction warrants further investigation.

Cell-type specific repertoires of HSPGs and their binding partners have been suggested to mediate the development of specific synaptic connections.^94^ We show that GPC1, which may act as a presynaptic GPR158 binding partner^26^, promotes the GPR158-PLCXD2 interaction, raising the intriguing possibility that SA content can be controlled via *trans*-synaptic interactions in an input-specific manner. Thus, SA incorporation could be facilitated at a select set of dendritic spines where a *trans*- synaptic HSPG-GPR158 interaction promotes binding of GPR158 to PLCXD2 to regulate its activity. HSPGs are not only expressed in neurons, but also in astrocytes^95^, which could additionally participate in modulating the GPR158-PLCXD2 association. Differential temporal expression patterns and HSPG identity are likely to be important factors determining which cellular source(s) of HSPGs are involved. Moreover, activity-dependent expression of HSPGs in both astrocytes^96^ and neurons^97^ provides an additional layer of temporal control over the GPR158-PLCXD2 interaction. We find that PLCXD2 localizes to a subset of dendritic spines. To what extent GPR158 and PLCXD2 are present together in the postsynaptic compartment and whether additional synaptic receptors exist to regulate PLCXD2’s activity remain outstanding questions that will be challenging to address given the lack of specific PLCXD2 antibodies.

Loss of the autism-associated gene *Klhl17*, an actin-binding protein that associates with SYNPO, interferes with postsynaptic SA localization and impairs dendritic spine maturation and plasticity.^16^ In addition, dysregulated synaptic SOCE has been implicated in various neurodegenerative diseases.^15^ Collectively, these studies highlight the importance of mechanisms controlling postsynaptic SA abundance and function for proper synapse development, underscoring the potential contribution of our findings to a better understanding of various brain disorders.

In conclusion, we identify a novel postsynaptic GPR158-PLCXD2 complex that controls SA abundance and the maturation of dendritic spines. Our data provide novel insight into GPR158-mediated signaling that likely has implications beyond its involvement in mood disorder.^27^ Finally, our findings shed new light on the molecular mechanisms via which synaptic cell-surface proteins instruct synapse development.

## Acknowledgments

We are grateful to Kirill A. Martemyanov and Thibaut Laboute for advice, discussion and sharing of unpublished findings. We thank Kirill A. Martemyanov, Franck Polleux, Pierre Vanderhaeghen and Marc Fivaz for critical reading of the manuscript, and members of the de Wit laboratory for helpful discussion and comments. We also thank Kristofer Davie (VIB Center for Brain & Disease Research Single-Cell Bioinformatics Expertise Unit) for bio-informatics support. We thank Jan Cools (VIB Center for Cancer Biology, Laboratory of Molecular Biology of Leukemia, Department of Human Genetics, KU Leuven, Leuven, Belgium) for sharing *H11^Cas^*^9^ mice and Vincent Bonin (Neuro-Electronics Research Flanders, Department of Biology, Leuven Brain Institute, KU Leuven, Leuven, Belgium) for sharing Rorb-IRES2-Cre mice. B.V. is supported by an FWO PhD fellowship 11A0419N. D.D. is supported by FWO Postdoctoral fellowships 12W5218N and 12W5221N, and an FWO Research Grant 1513320N.

A.C.N.F. is supported by an FWO Postdoctoral fellowship 1278323N. T.V. is supported by the Research Council of KU Leuven (C24M/21/028) and a Queen Elisabeth Medical Foundation for Neurosciences. J.T. is supported by a UGent Methusalem, ERC Advanced (CYRE, No. 340941) and an Interuniversity Attraction Poles (Belgian Science Policy; IAP-P7/13). J.d.W. is supported by FWO Project Grants G0C4518N, G0A8720N and G0A8320N; FWO EOS Grant G0H2818N; ERANET-NEURON TAO2PATHY; Queen Elisabeth Medical Foundation for Neurosciences and a KU Leuven Methusalem.

## Author contributions

Conceptualization: B.V., J.d.W.

Methodology: B.V., D.D., I.L., J.W., A.E.-A., J.T., T.V., S.M.

Investigation: B.V., L.A., J.V.ds., J.V.db., E.L., D.D., I.L., J.W., K.V., A.E.-A., A.C.N.F.

Funding acquisition: B.V. and J.d.W. Supervision: J.d.W.

Writing – original draft: B.V. and J.d.W.

Writing – review and editing: B.V. and J.d.W. with input from all authors

## Methods

### Animals

All animal experiments were conducted according to the KU Leuven ethical guidelines and approved by the KU Leuven Ethical Committee (approved protocol numbers ECD P183/2017 and P170/2021). Mice were maintained in a specific pathogen-free facility under standard housing conditions with continuous access to food and water. The health and welfare of the animals was supervised by a designated veterinarian. The KU Leuven animal facilities comply with all appropriate standards (cages, space per animal, temperature, light, humidity, food, water), and all cages are enriched with materials that allow the animals to exert their natural behavior. All mouse lines were maintained on a C57BL/6J background, bred in-house and raised in a temperature-controlled and humidity-controlled room with a 14–10 h light–dark cycle (lights on from 7:00 to 21:00). C57BL/6J mice (Charles River), *H11^Cas9^* (#027650) and *Rorb-IRES2- Cre* mice (#023526) were obtained from Jackson Laboratory. The *Gpr158* KO line (*Gpr158tm1(KOMP)Vlcg*) harboring a *LacZ* cassette was described previously.^26^ *Gpr158* and *Plcxd2* floxed mice were produced by Cyagen via homologous recombination, targeting exon 4 and exon 2 of the *Gpr158* and *Plcxd2* gene, respectively. L4-specific *Gpr158* cKO mice were generated by crossing *Gpr158^fl/+^:Rorb-Cre^+/-^* mice together to obtain *Gpr158^fl/fl^:Rorb-Cre* and *Gpr158^+/+^:Rorb-Cre* littermate controls. The same strategy was used to generate L4-specific *Plcxd2* cKO mice. To generate dcKO mice, *Gpr158^f/f^* and *Plcxd2^f/f^* mice were crossed to obtain *Gpr158^f/+^:Plcxd2^f/+^* mice. These mice were then crossed together to obtain *Gpr158^f/f^:Plcxd2^f/f^* and *Gpr158^w/w^:Plcxd2^w/w^* mice which were maintained as individual colonies. For euthanasia, animals were injected with an irreversible dose of ketamine– xylazine. Both males and females were used for all experiments. To the best of our knowledge, we are not aware of an influence of sex on the parameters analyzed in this study.

### Plasmids

Constructs were all generated using the Gibson Assembly Cloning Kit (New England Biolabs) by inserting the different DNA fragments (PCR-generated or gBlocks) in the final digested vectors. Full-length mouse GPR158 containing an N-terminal HA-tag or smURFP-tag were cloned into pcDNA3.1 (Thermo Fisher Scientific). Full-length cDNA encoding mouse PLCXD2 containing a C-terminal FLAG-tag in pcDNA3.1 (clone OMu06191, NM_001134480.1) was purchased from GenScript. The FLAG-tag was replaced with TdTomato to generate PLCXD2-TdTomato. GPR158-CFAST and PLCXD2-NFAST were produced by e-Zyvec, Polyplus. GPR158-CFAST contains full-length mouse GPR158 with an N-terminal HA-tag and a C-terminal CFAST11-tag. PLCXD2-NFAST contains full-length mouse PLCXD2 with a C-terminal FLAG-tag and NFAST-tag that are separated by a GS-linker. The catalytic dead mutant PLCXD2- FLAG CD where histidine residues at position 57 and 132 were replaced with leucines was generated by *in vitro* mutagenesis using the QuikChange II Site-Directed Mutagenesis Kit (Agilent) from PLCXD2-FLAG. pCAG_smFP HA (#59759), GFP-MAPPER (#117721), GFP-LiMETER-v2 (#113934), EGFP-ESyt2 (#66831) and GFP-C1-PLCdelta-PH (#21179) were obtained from Addgene. The following constructs were kindly provided: AcGFP-Sec61β (Franck Polleux, Columbia University Medical Center, USA), EGFP-Rab7 and EGFP-Rab11 (Casper Hoogenraad, Utrecht University, The Netherlands). PBOS-EGFP, pFUGW-mCherry and pFUGW-Cre-T2A- mCherry were described previously.^98^ PEGFP-GPR158 was described previously.^26^ All DNA constructs were verified by sequencing.

### Cell lines and transfections

HEK293T human embryonic kidney cells (available source material information: foetus) were obtained from American Type Culture Collection (ATCC) Cat #CRL- 11268. HEK293T cells were grown in DMEM (Thermo Fisher Scientific) supplemented with 10% fetal bovine serum (FBS, Thermo Fisher Scientific) and 1% penicillin/streptomycin (Thermo Fisher Scientific), and maintained in a humidified incubator of 5% CO2 at 37°C.

HEK239T cells were transfected using Fugene6 (Promega) with a DNA-to-Fugene ratio of 1:3 for 18-24 hours.

### Primary neuronal cultures and transfections

Neurons were cultured from E18 to E19 C57BL/6J WT mice embryos (hippocampal and cortical), from E18 to E19 to *H11^Cas9^* mice embryos (cortical) and from *Gpr158^fl/fl^* mice embryos (cortical). Dissected hippocampi and cortices were incubated with trypsin (0.25%, 15 min, 37°C; Thermo Fisher Scientific) in HBSS (Thermo Fisher Scientific) supplemented with 10mM HEPES (Thermo Fisher Scientific). After trypsin incubation, hippocampi and cortices were washed with MEM (Thermo Fisher Scientific) supplemented with 10% horse serum (Thermo Fisher Scientific) and 0.6% glucose (MEM-horse serum medium) 3 times. The cells were mechanically dissociated by repeatedly pipetting the tissue up and down in a flame-polished Pasteur pipette and then plated on poly-D-lysine (PDL, Millipore) and laminin (Sigma-Aldrich) coated glass coverslips (Glaswarenfabrik Karl Hecht) in 12-well plates, containing MEM-horse serum medium. Once neurons attached to the substrate, after 2 to 4 hours, the coverslips were flipped over an astroglial feeder layer in 12-well plates containing neuronal culture medium: Neurobasal medium (Thermo Fisher Scientific) supplemented with B27 (1:50, Thermo Fisher Scientific), 12mM glucose, glutamax (1:400, Thermo Fisher Scientific), penicillin/streptomycin (1:500, Thermo Fisher Scientific) and 25μM β-mercaptoethanol (Sigma-Aldrich). Neurons grew face down over the feeder layer but were kept separated from the glia. To prevent overgrowth of glia, neuron cultures were treated with 10μM 5-Fluoro-2′-deoxyuridine (Sigma-Aldrich) after 3 days. Cultures were maintained in a humidified incubator of 5% CO2 at 37°C.

Mouse cortical neurons from C57BL/6J and *Cas9^H^*^11^ mice were transfected at DIV12 using calcium phosphate with a total of 1µg DNA of the indicated constructs. Plasmids were diluted in Tris-EDTA buffer (10mM Tris-HCl and 2.5mM EDTA [pH 7.3]), followed by dropwise addition of CaCl2 solution (2.5M CaCl2 in 10mM HEPES [pH 7.2]) to the plasmid DNA-containing solution to give a final concentration of 250mM CaCl2. This solution was subsequently added to an equal volume of HEPES-buffered solution (274mM NaCl, 10mM KCl, 1.4mM Na2HPO4, 42mM HEPES [pH 7.2]) and vortexed gently for 3s. This mixture, containing precipitated DNA, was then added dropwise to the coverslips in a 12-well plate, containing 250μL of conditioned neuronal culture medium with 2mM kynurenic acid, followed by 2 hours incubation in a 37°C, 5% CO2 incubator. After 2 hours, the transfection solution was removed, after which 1 mL of conditioned neuronal culture medium with 2mM kynurenic acid slightly acidified with HCl (approximately 5 mM final concentration) was added to each coverslip, and the plate was returned to a 37°C, 5% CO2 incubator for 20 min. Finally, coverslips were then transferred to their original 12-well plate containing conditioned culture medium.

Mouse hippocampal neurons from C57BL/6J mice were transfected at DIV14 using Lipofectamine2000 (Thermo Fisher Scientific) with a total of 1µg DNA of the indicated constructs and 2µL Lipofectamine in Neurobasal medium. The DNA:lipofectamine mixture was added to the coverslips in a 12-well plate for 45 min before returning them to their original 12-well plate containing conditioned culture medium.

### MAPPIT

The cytoplasmic domain of human GPR158 containing amino acids 590-1136 was amplified and fused to the EpoR-LepR-F3 chimeric receptor in the pSEL (+2L) vector via SalI-NotI restriction cloning and screened against the human ORFeome v8.1 and ORFeome Collaboration prey library in the pMG1 vector as described previously.^99^

### Live cell imaging

For splitFAST experiments, HEK293T cells were cultured in 96-well plates (Greiner Bio-One) and transfected with the indicated constructs using Fugene6. After 24 hours, cells were treated with 2U/mL heparinase I/III (Sigma-Aldrich) or vehicle (20mM Tris-HCl [pH 7.5], 0.1mg/ml BSA, 4mM CaCl2) for 2 hours at 37°C. Following washes, cells were then incubated with 15 μg/mL recombinant mouse GPC1 (R&D Systems) in DMEM supplemented with 20mM HEPES for 1 hour at 37°C. After washing, medium was replaced with HBSS++ (Thermo Fisher Scientific) supplemented with 20mM HEPES containing 10μM HMBR (The Twinkle Factory) and cells were imaged live at 37°C. Primary cultured hippocampal neurons were transfected at DIV12 with the indicated constructs using Lipofectamine2000 as described above. Forty-eight hours later, live imaging was performed at 37°C in medium (119mM NaCl, 5mM KCl, 2mM CaCl2, 2mM MgCl2, 30mM glucose, and 10mM HEPES [pH 7.4]) containing 5μM HMBR.

For LiMETER experiments, HEK293T cells were cultured on glass coverslip-containing 35mm dishes (MatTek) and transfected with the indicated constructs using Fugene6. After 18-24 hours, cells were treated with 2U/mL heparinase I/III or vehicle (20mM Tris-HCl [pH 7.5], 0.1mg/ml BSA, 4mM CaCl2) for 2 hours at 37°C. Four hours prior to imaging, 1μM biliverdin dimethylester (Santa Cruz Biotechnology) – a cofactor for the far-red fluorescent protein smURFP^100^, was added to the cells. After washing, medium was replaced with HBSS++ supplemented with 20mM HEPES and cells were imaged live at 37°C.

### Immunocytochemistry and immunohistochemistry

#### Immunocytochemistry

HEK293T cells and primary cultured neurons were fixed in 4% paraformaldehyde (PFA)/4% sucrose in PBS for 15 min and permeabilized with 0.25% Triton X-100 for 5 min. Cells were then blocked in 10% bovine serum albumin (BSA) in PBS for 1 hour and incubated with primary antibodies diluted in 3% BSA in PBS overnight at 4°C. Following several washes, appropriate secondary antibodies were added for 1 hour at room temperature (RT). Coverslips were mounted using Prolong Gold Antifade (Thermo Fisher Scientific). Primary antibodies were the following: chicken anti-GFP (1:1000, Aves Labs), rabbit anti-HA (1:1000, Cell Signaling Technology), mouse anti-FLAG (1:1000, Cell Signaling Technology), rabbit anti-FLAG (1:1000, Cell Signaling Technology), rabbit anti-synaptopodin SE19 (1:500, Sigma-Aldrich) and mouse anti- PSD95 (1:250, Thermo Fisher Scientific). Fluorophore-conjugated secondary antibodies were from Jackson ImmunoResearch or Invitrogen. Alexa 647-conjugated phalloidin was acquired from Cell Signaling Technologies.

For surface immunolabeling of HA-GPR158, live HEK239T cells were incubated with rabbit anti-HA (1:500, Cell Signaling Technology) diluted in HEK239T cell medium for 20 min at RT. Cells were then fixed in 4% PFA/4% sucrose in PBS for 15 min at RT, blocked in 3% BSA in PBS for 1 hour and incubated with fluorophore-conjugated secondary antibodies in 3% BSA in PBS for 1 hour at RT. Following permeabilization, neurons were processed for immunocytochemistry as described above.

#### Immunohistochemistry

P14-P21 mice were anesthetized with ketamine (0.2 mg/gram body weight) and xylazine (0.02 mg/gram body weight) intraperitoneally administered, and transcardially perfused with 4% PFA in PBS. Brains were post-fixed overnight with 4% PFA in PBS at 4°C. For the analysis of dendritic spines, brains were perfused with 4% PFA (EM grade, Science Services), 2% sucrose diluted in 0.1M PB buffer pH 7.4 and post-fixed for 2 hours at RT. After washing in PBS, brains were embedded in 4% agarose and 50µm thick coronal sections were cut using a vibratome (Campden Instruments 7000smz). Heat-induced antigen retrieval was performed to immunolabel GPR158 by heating the sections for 30 min in a 10mM sodium citrate buffer (pH 6.0) containing 0.05% Tween20. Sections were permeabilized and blocked overnight at 4°C with 0.5% Triton x-100, 10% normal horse serum (NHS) and 0.5M glycine diluted in PBS-0.2% gelatin. Primary antibodies were incubated for 48 hours at 4°C. Following several washes, appropriate secondary antibodies were incubated for 24 hours at 4°C. Primary and secondary antibodies were diluted in 5% NHS and 0.5% triton X-100 in PBS-0.2% gelatin. Primary antibodies were the following: chicken anti-GFP (1:1000, Aves Labs), rat anti-HA (1:250, Roche), mouse anti-β-galactosidase (1:1000, Roche), rabbit anti- GPR158 (1:200, Sigma-Aldrich), rabbit anti-synaptopodin SE19 (1:500, Sigma-Aldrich), camelid PSD95-FluoTag-X2 (1:500, Synaptic Systems), guinea-pig anti- VGluT1 (1:000, Synaptic Systems), and guinea-pig anti-VGluT2 (1:1000, Synaptic Systems). Fluorophore-conjugated secondary antibodies were from Jackson ImmunoResearch or Invitrogen. Sections were mounted using Mowiol (Sigma-Aldrich).

#### RNAScope

C57BL/6J WT, *Plcxd2^fl/fl^:Rorb-Cre* and *Plcxd2^+/+^:Rorb-Cre* mice were transcardially perfused as described before. Fifty micrometers thick vibratome sections were mounted on SuperFrost Ultra Plus adhesion slides (Thermo Fisher Scientific) and warmed at 40°C for 15 minutes to improve adhesion. Then, sections were treated with H2O2 for 10 min and with protease IV for 20 min at RT. Hybridization was performed using the RNAscope™ Multiplex Fluorescent Detection Kit v2 (Advanced Cell Diagnostics) following manufacturer protocols. Briefly, *Rorb*, *Gpr158* and *Plcxd2* RNAScope probes were hybridized for 2 hours at 40°C followed by amplification steps and fluorescent labeling using OPAL dyes. To immunolabel for VGluT2, sections were next blocked for 1 hour at RT with 0.5% Triton X-100 and 5% NHS diluted in PBS. Primary antibody diluted in blocking solution was then added overnight at 4°C followed by secondary antibody incubation for 2 hours at RT. DAPI was used as nuclear stain. Sections were mounted using Mowiol. Imaging was performed using a ZEISS LSM880 in confocal mode equipped with a 63x Plan Apochromat (1.40) oil objective.

### Biochemistry

#### Co-immunoprecipitation

HEK293T cells were cultured in 6-well plates and transfected for 48 hours with the indicated constructs using Fugene6. After washing with cold PBS, cells were incubated with extraction buffer containing 150mM NaCl, 1% CHAPS (VWR) and 50mM HEPES (pH 7.4) supplemented with protease inhibitors (Roche) for 45 minutes at 4°C while gently shaking. Lysates were then spun at 14,800 rpm for 20 min at 4°C. Supernatants were precleared by adding 50µL Protein-G agarose beads (Thermo Fisher Scientific) for 1 hour at 4°C while rotating end-over-end. To pull down HA-GPR158, supernatants were incubated with 30µL anti-HA magnetic beads (Thermo Fisher Scientific) overnight at 4°C while rotating end-over-end. Anti-HA magnetic beads were then washed 3 times following manufacturer protocols, followed by 1 wash with MilliQ water using a magnetic stand. To pull down PLCXD2-FLAG, 5µg mouse anti-FLAG was added to the supernatants and incubated overnight at 4°C while rotating end-over-end. Then, supernatants were incubated with 30µL Protein-G agarose beads for 1 hour at 4°C while rotating end-over-end followed by 3 washes with cold extraction buffer and 1 wash with PBS. Proteins were eluted by heating Protein-G agarose beads and anti-HA magnetic beads for 15 min at 42°C in 50μL 2X sample buffer and analyzed by western blotting using rabbit anti-HA (1:1000, Cell Signaling Technology) and mouse anti-FLAG (1:1000, Sigma-Aldrich).

#### Transduction of neuronal cultures and collection of lysates

Primary cultured cortical neurons from *Gpr158^f/f^* mice were transduced at DIV2 with lentiviral vectors expressing Cre-T2A-mCherry or mCherry as control. Lysates were then harvested at DIV14 in Sodium Tris EDTA NP40 (STEN) buffer containing 50mM Tris HCl (pH 7.6), 150mM NaCl, 2mM EDTA, 0.2% NP-40, 1% Triton X-100 supplemented with protease inhibitors (Roche). Media was removed from each well and replaced with STEN buffer. After 30 min incubation while gently shaking at 4°C, lysates were collected and stored in tubes on ice for 10 min. Samples were then spun at 14,800 rpm for 10 min at 4°C and supernatant was collected. Protein concentrations were determined using the Bio-Rad protein assay kit, and GPR158 levels were analyzed by western blotting using rabbit anti-GPR158 (1:6000, kind gift Kirill Martemyanov) and rabbit anti-βIII-tubulin (1:5000, Abcam)

### Intracellular Ca^2+^-imaging

Between 18 and 24 hours after transfection with the indicated constructs, HEK239T cells were reseeded on PDL-coated glass coverslips to allow precise imaging of individual cells. Cells were incubated with 2µM Fura-2 AM (Thermo Fisher Scientific) for 40 min at 37°C, washed and mounted in an imaging chamber with standard bath solution containing 150mM NaCl, 6mM KCl, 2mM CaCl2, 1.5mM MgCl2, 10mM Glucose and 10mM HEPES (pH 7.4). Cells were left in standard bath solution for 10 min at RT before imaging. Fluorescence was measured during alternating illumination at 340 and 380 nm using an MT10 illuminator connected to an Olympus IX81 controlled by CellM software (Olympus). Changes in intracellular Ca^2+^ levels in individual cells were monitored as the ratio of the fluorescence at both excitation wavelengths (F340/F380) after correcting for background fluorescence, using custom-written routines in IgorPro 6.37. During imaging, cells were continuously perfused via a gravity-driven perfusion system, which ensured rapid (<2s) solution exchange around the imaged cells. To measure SOCE, cells were first perfused with a Ca^2+^-free solution containing 150mM NaCl, 6mM KCl, 1mM EGTA, 1.5mM MgCl2, 10mM Glucose and 10mM HEPES (pH 7.4), supplemented with the SERCA pump inhibitor cyclopiazonic acid (CPA; 20 µM) to cause passive store depletion. Reapplication of the standard, Ca^2+^-containing bath solution then resulted in SOCE. Transfected and non-transfected cells within a coverslip were identified based on GFP and TdTomato fluorescence. To reduce intrinsic variability between experiments, the store-operated calcium responses recorded from a coverslip were normalized to the mean response of the non-transfected cells within that experiment. Both the amplitude (ΔF340/F380) and the peak rate of rise (d(F340/F380)/dt) were quantified.

### Endogenous tagging

To endogenously tag PLCXD2, we modified a protocol based on the HiUGE (Homology-Independent Universal Gene Editing) technique.^39^ AAV plasmids containing a guide RNA (gRNA) cloning site downstream of an hU6 promoter, as well as the cassette to be inserted into genes of interest, were a kind gift from Scott Soderling. These plasmids were modified to express 2 gRNAs (targeting the gene of interest and the excisable cassette) as well as an mCherry reporter using the Gibson Assembly Cloning Kit. Furthermore, the initial 3X HA-tag was substituted with a smFP-HA tag amplified from pCAG_smFP HA.^101^ We selected 6 potential C-terminal insertion sites for HiUGE using Benchling. Six gRNAs corresponding to these sites were cloned into our plasmid according to previously published annealing and ligation protocols.^39^ We produced low-titer AAV supernatant for these 6 targets using a fast protocol. Briefly, single wells in a 12 well plate were plated with HEK cells and transfected with polyethyleneimine at ∼80% confluency with the 3 following plasmids: the pAdΔF6 helper, the RepCap 2/1, and one of the 6 gRNA-mCherry HiUGE plasmids. AAV-containing supernatant was collected after 3 days, filtered through a 0.45µm filter, and subsequently concentrated using 0.4mL 100.000MW Amicon centrifugation columns (Sigma-Aldrich). After 3 washes with PBS, the concentrated virus was aliquoted and stored at -80°C. We screened for efficient guides by transducing primary neuronal cultures from *H11^Cas9^* mice and immunolabeling for HA. Using this approach, we identified one gRNA (AGAGGATCGAGCTCTGACTA, 7 amino acids truncated from its C-terminus) as suitable for endogenous tagging of PLCXD2. The respective plasmid was used to produce high-titer AAV particles for *in vivo* experiments as described below.

### AAV production

AAV-TRE-DIO-FLPo, AAV-TRE-fDIO-GFP-IRES-tTA, AAV-TRE-Cre, AAV-SYN-DIO-GFP-IRES-tTA and the endogenous tagging plasmids were used to produce high-titer AAVs in-house. HEK293T cells were seeded in DMEM containing 10% FBS. Transfection mix, containing PEI and OptiMEM (Thermo Fisher Scientific), pAdΔF6, RepCap 2/1 or RepCap 2/9 and vector plasmid of interest were added to the cells in DMEM containing 1% FBS. Following transfection, cells were incubated in DMEM containing 5% FBS for 3 days. Cells were then harvested, spun at 1.000 g for 10 min at 4°C and pellets were lysed in lysis buffer (150mM NaCl and 50mM Tris HCl [pH 8.5]). Lysates were further frozen and thawed three times, spun at 2.000 g for 5 min at 4°C and Benzonase (Sigma-Aldrich) added at a concentration of 50U/mL to supernatants for 30 min at 37°C. Lysates were spun at 5.000 g for 20 min at RT and loaded onto an OptiPrep iodixanol gradient (Sigma-Aldrich) (60%, 40%, 25%, 15%) followed by ultracentrifugation at 300.000 g for 100 min at 12°C. Next, purified AAVs were carefully collected with an 18-gauge needle (Beckman Coulter) from between the 40% and 60% layers, and desalted and concentrated by centrifugation at 5,000 g for 30 min at 20 °C in a prerinsed Amicon Ultra-15 filter (Millipore). AAVs were aliquoted and stored at -80°C. AAV purity was tested by silver staining using the ProteoSilver Silver Stain Kit (Sigma PROTSIL1-1KT).

### Neonatal stereotaxic injections

P0 *Gpr158^fl/fl^:Rorb-Cre*, *Gpr158^+/+^:Rorb-Cre*, *Plcxd2^fl/fl^:Rorb-Cre* and *Plcxd2^+/+^:Rorb- Cre* pups were injected with 50nL of a mixture containing AAV-TRE-DIO-FLPo (Addgene #118027) and AAV-TRE-fDIO-GFP-IRES-tTA (Addgene #118026) in a 1:50 ratio in the barrel cortex, directly through the skin and skull using a Nanoject III Auto-Nanoliter Injector (Drummond). P0 *Gpr158^+/+^:Plcxd2^+/+^*, *Gpr158fl/fl:Plcxd2^+/+^* and *Gpr158^fl/fl^:Plcxd2^fl/fl^* pups were injected with 50nL of a mixture containing AAV-TRE-Cre (Addgene #69136) and AAV-SYN-DIO-GFP-IRES-tTA (Addgene #85006) in a 1:50 ratio in the barrel cortex, directly through the skin and skull using a Nanoject III Auto-Nanoliter Injector (Drummond). P0 pups were anesthetized by hypothermia and then stabilized on a glass petri dish filled with ice to sustain anesthesia during injections. To endogenously tag PLCXD2 *in vivo*, P7 *H11^Cas9^* pups were anesthetized with 5% isoflurane and Duratears was applied to the eyes to prevent them from drying out. Mice were placed in a mouse stereotact (KOPF). During the rest of the procedure 2.5% isoflurane was constantly administered. After shaving and disinfecting the mouse’s head, local anesthesia was administered by a subcutaneous injection with 50µl lidocaine (xylocain 1%). An incision was made on the skin to reveal the skull and 250nL of AAV-PLCXD2-smHA was injected in the barrel cortex. Following injection, the incision was stitched with surgical glue (Millpledge Veterinary). After 6 hours, their health was examined and pups were injected with 0.1mg/kg buprenorphine.

### Microscopy and image analysis

Unless stated otherwise, all imaging was performed on a ZEISS LSM880 in Airyscan mode with a 63x Plan Apochromat (1.40) oil objective. Optimal settings suggested by the ZEN Black software were used to determine pixel size, zoom and optical sectioning. Raw Airyscan images were processed using the Airyscan processing tool set to default deconvolution strength. Tile scans of brain sections were taken on a ZEISS LSM880 in confocal mode with a 20x air objective. Across experiments, identical laser power and detector gain were used when comparing different conditions.

#### Structured illumination microscopy

HEK239T cells were cultured on chemically cleaned #1.5H High Performance coverslips (Marienfeld Superior) and transfected with the indicated constructs using Fugene6. Twenty-four hours later, cells were immunolabeled as described above and mounted using Prolong Diamond Antifade (Thermo Fisher Scientific). Imaging was performed on a Nikon Ti2 N-SIM S equipped with a 100x SR HP Apo TIRF (1.49) oil objective and an ORCA-Flash 4.0 (C13440, Hamamatsu) camera. The setup was controlled by NIS Elements 5.30.07 (Build 1569). Images were acquired in TIRF-SIM mode. Reconstruction was done using NIS slice reconstruction with optimized parameters for each channel. Images were analyzed in batch with NIS-Elements 5.42.03 (Nikon). The GA3 analysis protocol included background subtraction of a constant (noise), manual thresholding and filtering of small objects. Parent-child analysis on the binary masks was performed to find the nearest neighbor distances of PLCXD2 to MAPPER. To obtain random data, we used the same set of images, but regions of interests (ROIs) of one channel were paired with non-corresponding ROIs of the other channel. We paired each ROI of MAPPER from 5 different cells with non-corresponding ROIs of PLCXD2 from 5 other cells.

#### LiMETER

HEK239T cells were cultured on glass coverslip-containing 35mm dishes and transfected with the indicated constructs using Fugene6. Eighteen to 24 hours later cells were imaged on an inverted Nikon TiE microscope equipped with a 100x TIRF (1.49) oil objective, coupled to a SAFe360 module (Abbelight) and two ORCA-Fusion Digital CMOS cameras (C14440-20P, Hamamatsu) for simultaneous acquisition. The setup was controlled by NEO software (Abbelight). Eight-hundred frames were recorded at 100 msec/frame. Cells were assessed for TdTomato and smURFP fluorescence before TIRF time-lapse imaging of LiMETER. Images were analyzed in batch with NIS-Elements 5.42.03 (Nikon). The GA3 analysis protocol included rolling ball background subtraction with a 5µm radius followed by a bright spot detection with a 6µm diameter and manual intensity thresholding. Throughout the time-lapse, the sum of the intensity of identified puncta was analyzed and normalized by cell area.

#### Plasma membrane PIP2 levels

HEK239T cells transfected with the indicated constructs were imaged near their adherent surface to allow clear visualization of PM and cytosolic compartments. Following automated thresholding in Fiji (ImageJ), integrated intensity of PH-PLCδ- GFP fluorescence was measured in ROIs encompassing the PM and the cytosol, and normalized to the corresponding ROI area. The PH-PLCδ-GFP PM/cytosol ratio was then calculated by dividing these values.

#### Analysis of spinous F-actin and SYNPO

Primary cultured hippocampal neurons were transfected with the indicated constructs using Lipofectamine2000. Forty-eight hours later, neurons were immunolabeled as described above. PLCXD2-FLAG and F-actin or SYNPO fluorescence were thresholded in Fiji (imageJ), converted to binary masks and transferred to the GFP image from which the percentage of PLCXD2-positive spines containing SYNPO and vice versa were calculated. F-actin and SYNPO fluorescence were measured as the integrated intensity in PLCXD2-positive or PLCXD2-negative dendritic spines.

#### Dendritic spine imaging and analysis

P0 pups were injected with the indicated AAVs as described above. Following 2 weeks of expression, mice were perfused and brain sections were prepared and immunolabeled for GFP and SYNPO as described above. One to three secondary dendrites spanning 75µm-175µm each were imaged per neuron. Dendritic protrusions were reconstructed with Neurolucida360 (MBF Bioscience) and classified using standard morphological parameters in Neurolucida Explorer. Dendritic protrusions were classified as filopodia when no discernable spine head was detected. To analyze SYNPO-containing dendritic spines, SYNPO puncta were reconstructed with Neurolucida360 using the Puncta tool and their presence in dendritic spines was analyzed using Neurolucida Explorer. Identical puncta detection settings were used across conditions.

#### splitFAST

HEK239T cells cultured in 96-well plates were imaged on a ZEISS Celldiscoverer 7 in widefield mode equipped with a 20x (0.95) air objective. For each condition, (vehicle, heparinase I/III treated or heparinase I/III + GPC1 treated) at least 3 wells per experiment were imaged, where 3 ROIs of identical size were randomly chosen per well. Images were subjected to rolling ball background subtraction and FAST fluorescence intensity was measured as the integrated intensity following automated thresholding in Fiji (ImageJ).

### Statistics

All data are graphed as mean ± SEM. Statistical testing was performed using GraphPad Prism (GraphPad Software). For normally distributed data, as determined by D’Agostino and Pearson test, differences were analyzed using a two-tailed Student’s t-test and a one-way ANOVA test when comparing 3 experimental groups. The Mann–Whitney and Kruskal–Wallis tests were used when criteria for normality were not met. Bonferroni’s, Dunn’s and Tukey’s tests were used as multiple-comparison post hoc tests.

## Multimedia files

**Movie S1.** Time-lapse imaging of LiMETER in HEK239T cells co-expressing LiMETER together with TdTomato (left), with PLCXD2-TdTomato (middle) or with PLCXD2- TdTomato and smURFP-GPR158 (right). Images where acquired every 100 ms for 80 sec. Playback rate is 10 frames per second.

**Movie S2.** Time-lapse imaging of HEK239T cells expressing GFP-MAPPER and PLCXD2-TdTomato. Images where acquired every 10 sec for 10 min. Playback rate is 10 frames per second.

**Movie S3.** Time-lapse imaging of the boxed area in Movie S2. Images where acquired every 10 sec for 10 min. Playback rate is 10 frames per second.

**Movie S4.** Time-lapse imaging of HEK239T cells expressing GFP-MAPPER alone. Images where acquired every 10 sec for 10 min. Playback rate is 10 frames per second.

**Movie S5.** Time-lapse imaging of the boxed area in Movie S4. Images where acquired every 10 sec for 10 min. Playback rate is 10 frames per seconds.

## References

1. O’Rourke, N. A., Weiler, N. C., Micheva, K. D. & Smith, S. J. Deep molecular diversity of mammalian synapses: Why it matters and how to measure it. Nat. Rev. Neurosci. 13, 365–379 (2012).

2. Emes, R. D. & Grant, S. G. N. Evolution of Synapse Complexity and Diversity. Annu. Rev. Neurosci. 35, 111–131 (2012).

3. Grant, S. G. N. Synapse diversity and synaptome architecture in human genetic disorders. Hum. Mol. Genet. 28, R219–R225 (2019).

4. Wu, Y. et al. Contacts between the endoplasmic reticulum and other membranes in neurons. Proc. Natl. Acad. Sci. U. S. A. 114, E4859–E4867 (2017).

5. Spacek, J. & Harris, K. M. Three-dimensional organization of smooth endoplasmic reticulum in hippocampal CA1 dendrites and dendritic spines of the immature and mature rat. J. Neurosci. Off. J. Soc. Neurosci. 17, 190–203 (1997).

6. Deller, T., Merten, T., Roth, S. U., Mundel, P. & Frotscher, M. Actin-associated protein synaptopodin in the rat hippocampal formation: Localization in the spine neck and close association with the spine apparatus of principal neurons. J. Comp. Neurol. 418, 164–181 (2000).

7. Orth, C. B. et al. Lamina-specific distribution of synaptopodin, an actin-associated molecule essential for the spine apparatus, in identified principal cell dendrites of the mouse hippocampus. J. Comp. Neurol. 487, 227–239 (2005).

8. Deller, T. et al. Synaptopodin-deficient mice lack a spine apparatus and show deficits in synaptic plasticity. Proc. Natl. Acad. Sci. U. S. A. 100, 10494–10499 (2003).

9. Jedlicka, P. et al. Impairment of in vivo theta-burst long-term potentiation and network excitability in the dentate gyrus of synaptopodin-deficient mice lacking the spine apparatus and the cisternal organelle. Hippocampus 19, 130–140 (2009).

10. Zhang, X. L. et al. Essential role for synaptopodin in dendritic spine plasticity of the developing hippocampus. J. Neurosci. 33, 12510–12518 (2013).

11. Dubes, S. et al. miR -124-dependent tagging of synapses by synaptopodin enables input-specific homeostatic plasticity . EMBO J. 41, 1–26 (2022).

12. Lenz, M. et al. All-trans retinoic acid induces synaptic plasticity in human cortical neurons. Elife 10, 1–20 (2021).

13. Yap, K. et al. The actin-modulating protein synaptopodin mediates long-term survival of dendritic spines. Elife 9, 1–31 (2020).

14. Maggio, N. & Vlachos, A. Synaptic plasticity at the interface of health and disease: New insights on the role of endoplasmic reticulum intracellular calcium stores. Neuroscience 281, 135–146 (2014).

15. Zhang, I. & Hu, H. Store-Operated Calcium Channels in Physiological and Pathological States of the Nervous System. Front. Cell. Neurosci. 14, 600758 (2020).

16. Hu, H. T. et al. Autism-related KLHL17 and SYNPO act in concert to control activity-dependent dendritic spine enlargement and the spine apparatus. PLoS Biol. 21, 1–30 (2023).

17. Perez-Alvarez, A. et al. Endoplasmic reticulum visits highly active spines and prevents runaway potentiation of synapses. Nat. Commun. 11, 1–10 (2020).

18. Konietzny, A. et al. Myosin V regulates synaptopodin clustering and localization in the dendrites of hippocampal neurons. J. Cell Sci. 132, (2019).

19. Konietzny, A. et al. Caldendrin and myosin V regulate synaptic spine apparatus localization via ER stabilization in dendritic spines. EMBO J. 41, 1–24 (2022).

20. Ng, A. N. & Toresson, H. γ-Secretase and metalloproteinase activity regulate the distribution of endoplasmic reticulum to hippocampal neuron dendritic spines. FASEB J. 22, 2832–2842 (2008).

21. Velikanov, G. A. & De Camilli, P. Endoplasmic reticulum-plasma membrane contact sites. Annu. Rev. Biochem. 55, 445–451 (2017).

22. Spacek, J. Relationships between synaptic junctions, puncta adhaerentia and the spine apparatus at neocortical axo-spinous synapses. A serial section study. Anat. Embryol. (Berl). 173, 129–135 (1985).

23. Spacer, J. & Harris, K. M. Three-dimensional organization of cell adhesion junctions at synapses and dendritic spines in area CA1 of the rat hippocampus. J. Comp. Neurol. 393, 58–68 (1998).

24. Falahati, H., Wu, Y., Feuerer, V., Simon, H.-G. & De Camilli, P. Proximity proteomics of synaptopodin provides insight into the molecular composition of the spine apparatus of dendritic spines. Proc. Natl. Acad. Sci. U. S. A. 119, e2203750119 (2022).

25. Chen, Y. J., Quintanilla, C. G. & Liou, J. Recent insights into mammalian ER– PM junctions. Curr. Opin. Cell Biol. 57, 99–105 (2019).

26. Condomitti, G. et al. An Input-Specific Orphan Receptor GPR158-HSPG Interaction Organizes Hippocampal Mossy Fiber-CA3 Synapses. Neuron 100, 201–215.e9 (2018).

27. Sutton, L. P. et al. Orphan receptor GPR158 controls stress-induced depression. Elife 7, 1–27 (2018).

28. Khrimian, L. et al. Gpr158 mediates osteocalcin’s regulation of cognition. J. Exp. Med. 214, 2859–2873 (2017).

29. Orlandi, C. et al. Transsynaptic Binding of Orphan Receptor GPR179 to Dystroglycan-Pikachurin Complex Is Essential for the Synaptic Organization of Photoreceptors. Cell Rep. 25, 130–145.e5 (2018).

30. Laboute, T. et al. Orphan receptor GPR158 serves as a metabotropic glycine receptor: mGlyR. Science (80-. ). 379, 1352–1358 (2023).

31. Orlandi, C. et al. Orphan receptor GPR158 is an allosteric modulator of RGS7 catalytic activity with an essential role in dictating its expression and localization in the brain. J. Biol. Chem. 290, 13622–13639 (2015).

32. Patil, D. N. et al. Cryo-EM structure of human GPR158 receptor coupled to the RGS7-Gβ5 signaling complex. Science 375, 86–91 (2022).

33. Lievens, S. et al. Proteome-scale binary interactomics in human cells. Mol. Cell. Proteomics 15, 3624–3639 (2016).

34. Belgard, T. G. et al. A transcriptomic atlas of mouse neocortical layers. Neuron 71, 605–616 (2011).

35. Li, H. et al. Laminar and Columnar Development of Barrel Cortex Relies on Thalamocortical Neurotransmission. Neuron 79, 970–986 (2013).

36. Klingler, E. et al. A Translaminar Genetic Logic for the Circuit Identity of Intracortically Projecting Neurons. Curr. Biol. 332–339 (2019). doi:10.1016/j.cub.2018.11.071

37. Zeisel, A. et al. Molecular Architecture of the Mouse Nervous System. Cell 174, 999–1014.e22 (2018).

38. Tebo, A. G. & Gautier, A. A split fluorescent reporter with rapid and reversible complementation. Nat. Commun. 10, 2822 (2019).

39. Gao, Y. et al. Plug-and-Play Protein Modification Using Homology-Independent Universal Genome Engineering. Neuron 103, 583–597.e8 (2019).

40. Gellatly, S. A., Kalujnaia, S. & Cramb, G. Cloning, tissue distribution and sub- cellular localisation of phospholipase C X-domain containing protein (PLCXD) isoforms. Biochem. Biophys. Res. Commun. 424, 651–656 (2012).

41. Mondin, V. E. et al. PTEN reduces endosomal PtdIns(4,5)P2 in a phosphatase-independent manner via a PLC pathway. J. Cell Biol. 218, jcb.201805155 (2019).

42. Chang, C. L. et al. Feedback regulation of receptor-induced ca2+ signaling mediated by e-syt1 and nir2 at endoplasmic reticulum-plasma membrane junctions. Cell Rep. 5, 813–825 (2013).

43. Giordano, F. et al. PI(4,5)P(2)-dependent and Ca(2+)-regulated ER-PM interactions mediated by the extended synaptotagmins. Cell 153, 1494–1509 (2013).

44. Liou, J., Fivaz, M., Inoue, T. & Meyer, T. Live-cell imaging reveals sequential oligomerization and local plasma membrane targeting of stromal interaction molecule 1 after Ca2+ store depletion. Proc. Natl. Acad. Sci. U. S. A. 104, 9301–9306 (2007).

45. Park, C. Y. et al. STIM1 Clusters and Activates CRAC Channels via Direct Binding of a Cytosolic Domain to Orai1. Cell 136, 876–890 (2009).

46. Walsh, C. M. et al. Role of phosphoinositides in STIM1 dynamics and store- operated calcium entry. Biochem. J. 425, 159–168 (2010).

47. Calloway, N. et al. Stimulated association of STIM1 and Orai1 is regulated by the balance of PtdIns(4,5)P 2 between distinct membrane pools. J. Cell Sci. 124, 2602–2610 (2011).

48. Bhardwaj, R., Müller, H. M., Nickel, W. & Seedorf, M. Oligomerization and Ca2+ /calmodulin control binding of the ER Ca2+ -sensors STIM1 and STIM2 to plasma membrane lipids. Biosci. Rep. 33, (2013).

49. Cohen, H. A., Zomot, E., Nataniel, T., Militsin, R. & Palty, R. The SOAR of STIM1 interacts with plasma membrane lipids to form ER-PM contact sites. Cell Rep. 42, 112238 (2023).

50. He, L. et al. Optical control of membrane tethering and interorganellar communication at nanoscales. Chem. Sci. 8, 5275–5281 (2017).

51. Prakriya, M. & Lewis, R. S. Store-Operated Calcium Channels. Physiol. Rev. 95, 1383–1436 (2015).

52. Zhang, S. L. et al. STIM1 is a Ca2+ sensor that activates CRAC channels and migrates from the Ca2+ store to the plasma membrane. Nature 437, 902–905 (2005).

53. Lewis, R. S. The molecular choreography of a store-operated calcium channel. Nature 446, 284–287 (2007).

54. Saarikangas, J., Zhao, H. & Lappalainen, P. Regulation of the actin cytoskeleton-plasma membrane interplay by phosphoinositides. Physiol. Rev. 90, 259–289 (2010).

55. Korkotian, E., Frotscher, M. & Segal, M. Synaptopodin regulates spine plasticity: mediation by calcium stores. J. Neurosci. Off. J. Soc. Neurosci. 34, 11641–11651 (2014).

56. Harris, J. A. et al. Anatomical characterization of Cre driver mice for neural circuit mapping and manipulation. Front. Neural Circuits 8, 1–16 (2014).

57. Lin, R. et al. Cell-type-specific and projection-specific brain-wide reconstruction of single neurons. Nat. Methods 15, 1033–1036 (2018).

58. Mizuno, H. et al. NMDAR-regulated dynamics of layer 4 neuronal dendrites during thalamocortical reorganization in neonates. Neuron 82, 365–379 (2014).

59. Luo, W. et al. Supernova: A Versatile Vector System for Single-Cell Labeling and Gene Function Studies in vivo. Sci. Rep. 6, 35747 (2016).

60. Geiger, T., Wehner, A., Schaab, C., Cox, J. & Mann, M. Comparative proteomic analysis of eleven common cell lines reveals ubiquitous but varying expression of most proteins. Mol. Cell. Proteomics 11, M111.014050 (2012).

61. Kadamur, G. & Ross, E. M. Mammalian phospholipase C. Annu. Rev. Physiol. 75, 127–154 (2013).

62. Katan, M. & Cockcroft, S. Phospholipase C families: Common themes and versatility in physiology and pathology. Prog. Lipid Res. 80, 101065 (2020).

63. Matt, L., Kim, K., Chowdhury, D. & Hell, J. W. Role of palmitoylation of postsynaptic proteins in promoting synaptic plasticity. Front. Mol. Neurosci. 12, 1–19 (2019).

64. Li, S. et al. pCysMod: Prediction of Multiple Cysteine Modifications Based on Deep Learning Framework. Front. Cell Dev. Biol. 9, 1–10 (2021).

65. Hicks, S. N. et al. General and Versatile Autoinhibition of PLC Isozymes. Mol. Cell 31, 383–394 (2008).

66. Lyon, A. M., Begley, J. A., Manett, T. D. & Tesmer, J. J. G. Molecular mechanisms of phospholipase C β3 autoinhibition. Structure 22, 1844–1854 (2014).

67. Wang, X. et al. Three-dimensional intact-tissue sequencing of single-cell transcriptional states. Science (80-. ). 361, (2018).

68. Jha, A. et al. Anoctamin 8 tethers endoplasmic reticulum and plasma membrane for assembly of Ca(2+) signaling complexes at the ER/PM compartment. EMBO J. 38, (2019).

69. Besprozvannaya, M. et al. GRAM domain proteins specialize functionally distinct ER-PM contact sites in human cells. Elife 7, 1–25 (2018).

70. Takeshima, H., Hoshijima, M. & Song, L. S. Ca2+ microdomains organized by junctophilins. Cell Calcium 58, 349–356 (2015).

71. Ghai, R. et al. ORP5 and ORP8 bind phosphatidylinositol-4, 5-biphosphate (PtdIns(4,5)P2) and regulate its level at the plasma membrane. Nat. Commun.8, (2017).

72. Wang, H. et al. ORP2 Delivers Cholesterol to the Plasma Membrane in Exchange for Phosphatidylinositol 4, 5-Bisphosphate (PI(4,5)P 2 ). Mol. Cell 73, 458-473.e7 (2019).

73. Weber-Boyvat, M. et al. ORP/Osh mediate cross-talk between ER-plasma membrane contact site components and plasma membrane SNAREs. Cell. Mol. Life Sci. 78, 1689–1708 (2021).

74. Lees, J. A. et al. Lipid transport by TMEM24 at ER-plasma membrane contacts regulates pulsatile insulin secretion. Science (80-. ). 355, (2017).

75. Sun, E. W. et al. Lipid transporter TMEM24/C2CD2L is a Ca2+-regulated component of ER–plasma membrane contacts in mammalian neurons. Proc. Natl. Acad. Sci. U. S. A. 116, 5775–5784 (2019).

76. Kirmiz, M. et al. Remodeling neuronal ER-PM junctions is a conserved nonconducting function of Kv2 plasma membrane ion channels. Mol. Biol. Cell 29, 2410–2432 (2018).

77. Kirmiz, M., Vierra, N. C., Palacio, S. & Trimmer, J. S. Identification of VAPA and VAPB as Kv2 channel-interacting proteins defining endoplasmic reticulum– plasma membrane junctions in mammalian brain neurons. J. Neurosci. 38, 7562–7584 (2018).

78. Sahu, G. et al. Junctophilin Proteins Tether a Cav1-RyR2-KCa3.1 Tripartite Complex to Regulate Neuronal Excitability. Cell Rep. 28, 2427–2442.e6 (2019).

79. Vierra, N. C., Kirmiz, M., List, D. van der, Santana, L. F. & Trimmer, J. S. Kv2.1 mediates spatial and functional coupling of L-type calcium channels and ryanodine receptors in neurons. Elife 702514 (2019). doi:10.1101/702514

80. Perni, S. & Beam, K. G. Neuronal junctophilins recruit specific Cav and RyR isoforms to ER-PM junctions and functionally alter Cav2.1 and Cav2.2. Elife 10, 1–31 (2021).

81. Kikuma, K., Li, X., Kim, D., Sutter, D. & Dickman, D. K. Extended synaptotagmin localizes to presynaptic ER and promotes neurotransmission and synaptic growth in drosophila. Genetics 207, 993–1006 (2017).

82. Gallo, A. et al. Role of the Sec22b–E-Syt complex in neurite growth and ramification. J. Cell Sci. 133, (2020).

83. Mao, R., Tong, C. & Liu, J. J. E-Syt1 Regulates Neuronal Activity-Dependent Endoplasmic Reticulum–Plasma Membrane Junctions and Surface Expression of AMPA Receptors. Contact 6, (2023).

84. Zhang, M., Li, X., Zhuo, S., Yang, M. & Yu, Z. Enriched Environment Enhances Sociability Through the Promotion of ESyt1-Related Synaptic Formation in the Medial Prefrontal Cortex. Mol. Neurobiol. (2023). doi:10.1007/s12035-023-03742-9

85. Courjaret, R., Prakriya, M. & Machaca, K. SOCE as a regulator of neuronal activity. J. Physiol. 0, 1–14 (2023).

86. Basnayake, K. et al. Nanoscale molecular architecture controls calcium diffusion and ER replenishment in dendritic spines. Sci. Adv. 7, (2021).

87. Sun, S. et al. Reduced synaptic STIM2 expression and impaired store-operated calcium entry cause destabilization of mature spines in mutant presenilin mice. Neuron 82, 79–93 (2014).

88. Zhang, H. et al. Neuronal store-operated calcium entry and mushroom spine loss in amyloid precursor protein knock-in mouse model of Alzheimer’s disease. J. Neurosci. 35, 13275–13286 (2015).

89. Yap, K. A. F. et al. STIM2 regulates AMPA receptor trafficking and plasticity at hippocampal synapses. Neurobiol. Learn. Mem. 138, 54–61 (2017).

90. Korkotian, E., Oni-Biton, E. & Segal, M. The role of the store-operated calcium entry channel Orai1 in cultured rat hippocampal synapse formation and plasticity. J. Physiol. 595, 125–140 (2017).

91. Maneshi, M. M. et al. Orai1 Channels Are Essential for Amplification of Glutamate-Evoked Ca2+ Signals in Dendritic Spines to Regulate Working and Associative Memory. Cell Rep. 33, 108464 (2020).

92. Garcia-Alvarez, G. et al. STIM2 regulates PKA-dependent phosphorylation and trafficking of AMPARs. Mol. Biol. Cell 26, 1141–1159 (2015).

93. Jeong, E., Kim, Y., Jeong, J. & Cho, Y. Structure of the class C orphan GPCRGPR158 in complex with RGS7-Gβ5. Nat. Commun. 12, 1–11 (2021).

94. Condomitti, G. & de Wit, J. Heparan Sulfate Proteoglycans as Emerging Players in Synaptic Specificity. Front. Mol. Neurosci. 11, 1–14 (2018).

95. Farhy-Tselnicker, I. et al. Astrocyte-Secreted Glypican 4 Regulates Release of Neuronal Pentraxin 1 from Axons to Induce Functional Synapse Formation. Neuron 96, 428–445.e13 (2017).

96. Farhy-Tselnicker, I. et al. Activity-dependent modulation of synapse-regulating genes in astrocytes. Elife 10, 1–43 (2021).

97. Butko, M. T. et al. In vivo quantitative proteomics of somatosensory cortical synapses shows which protein levels are modulated by sensory deprivation. Proc. Natl. Acad. Sci. 110, E726–E735 (2013).

98. Ribeiro, L. F. et al. SorCS1-mediated sorting in dendrites maintains neurexin axonal surface polarization required for synaptic function. PLoS Biol. 17, e3000466 (2019).

99. Lemmens, I., Lievens, S. & Tavernier, J. MAPPIT, a Mammalian Two-Hybrid Method for In-Cell Detection of Protein-Protein Interactions. Methods Mol. Biol. 1278, 447–455 (2015).

100. Rodriguez, E. A. et al. A far-red fluorescent protein evolved from a cyanobacterial phycobiliprotein. Nat. Methods 13, 763–769 (2016).

101. Viswanathan, S. et al. High-performance probes for light and electron microscopy. Nat. Methods 12, 568–576 (2015).

