## Supplemental information for "Cell-surface receptor-mediated regulation of synaptic organelle distribution controls dendritic spine maturation"

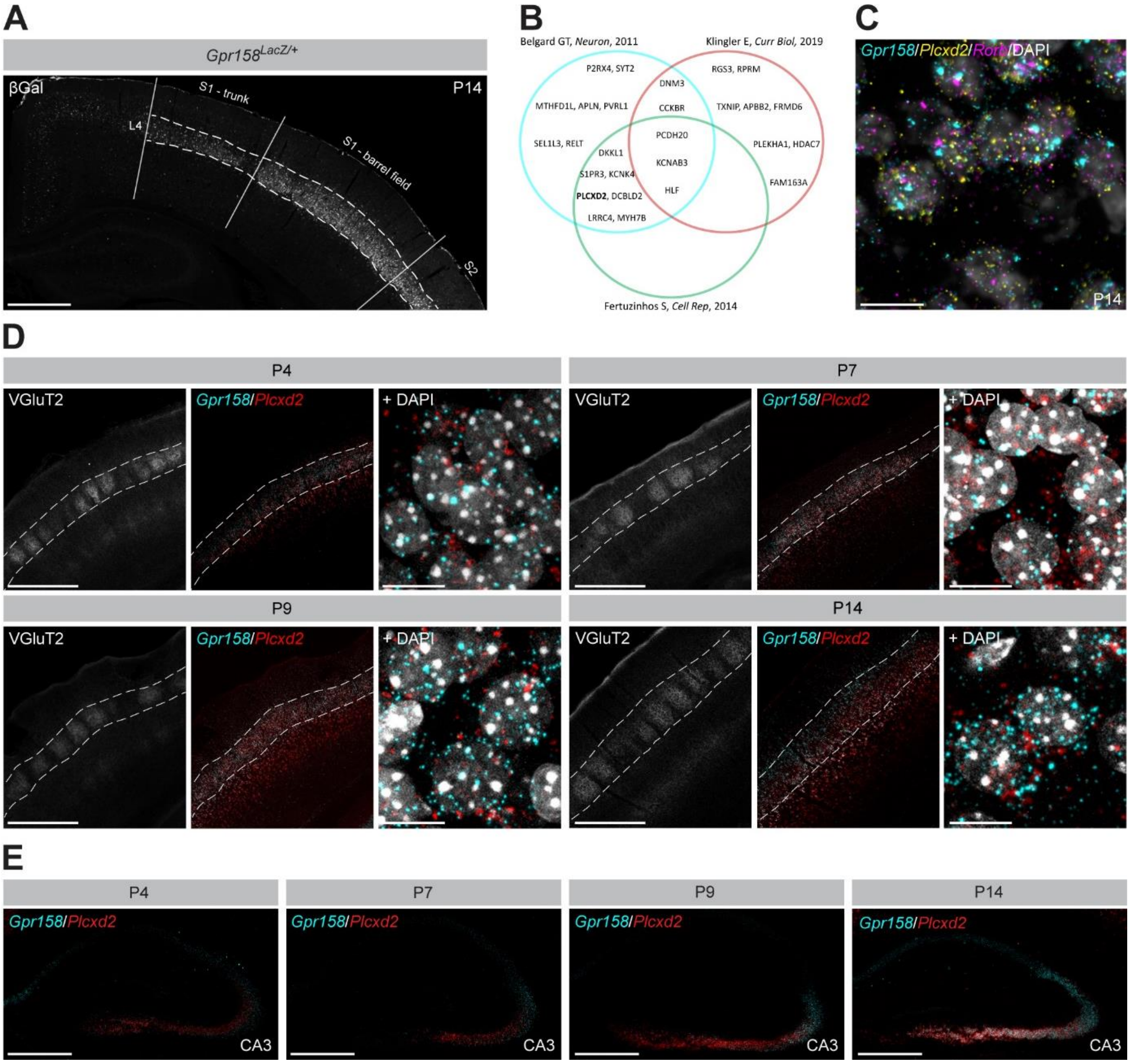


**Supplementary figure 1. Identification of a novel postsynaptic GPR158-PLCXD2 complex (see also Figure 1).**

**(A)** βGal immunohistochemistry demonstrates strong enrichment of *Gpr158* in L4 of the barrel cortex. VGluT2 immunohistochemistry (not shown) was used to delineate L4. Scale bar 500 µm.

**(B)** L4-enriched transcripts gathered from Belgard GT, *Neuron*, 2011 (cyan); Klingler E, Curr Biol, 2019 (red) and Fertuzinhos S, *Cell Rep*, 2014 (green) cross-referenced with putative GPR158 binding partners from the MAPPIT screen.

**(C)** P14 mouse coronal brain sections from the barrel cortex were probed for *Gpr158* (cyan), *Plcxd2* (yellow) and *Rorb* (magenta) using RNAScope at P14. Scale bar 10 µm.

**(D)** P4, P7, P9 and P14 mouse coronal brain sections from the barrel cortex were probed for *Gpr158* (cyan) and *Plcxd2* (red) using RNAScope, and immunostained for VGluT2 (gray) to delineate L4. Scale bar 500 µm, inset 10 µm.

**(E)** P4, P7, P9 and P14 mouse coronal brain sections from the hippocampus were probed for *Gpr158* (cyan) and *Plcxd2* (red) using RNAScope. Scale bar 500 µm.

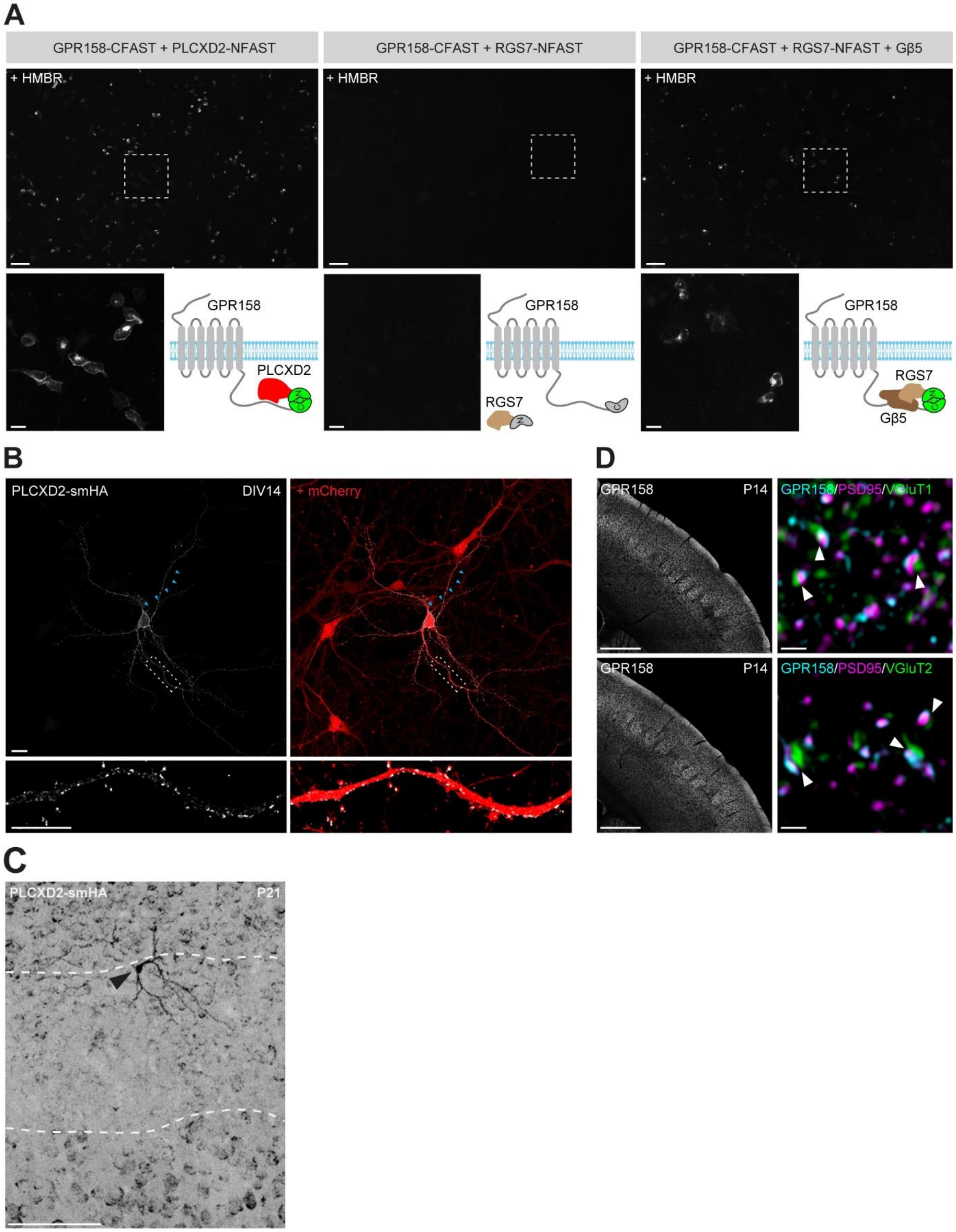


**Supplementary figure 2. Identification of a novel postsynaptic GPR158-PLCXD2 complex (see also Figure 1).**

**(A)** HEK239T cells co-expressing GPR158-CFAST and PLCXD2-NFAST (left), co-expressing GPR158-CFAST and RGS7-NFAST (middle) or co-expressing GPR158-

CFAST, RGS7-NFAST and Gβ5 (right) were visualized live in the presence of 10µM HMBR. Scale bar 100 µm, inset 20 µm.

**(B)** DIV14 cultured *H11*$^{Cas9}$ mouse cortical neuron demonstrating endogenously tagged PLCXD2 expression (gray), visualized by HA immunolabeling. The targeting cassette also expresses TdTomato (red) to visualize neuronal morphology. Blue arrowheads indicate the axon. Scale bar 20 µm, inset 10 µm.

**(C)** Overview image of an endogenously tagged PLCXD2-expressing neuron in L4 of the primary somatosensory cortex. High-magnification images in Fig. 1I are derived from this neuron.

**(D)** GPR158 (gray and cyan) immunoreactivity in a P14 mouse coronal brain section demonstrates a barrel-like pattern in L4 of the primary somatosensory cortex. High-magnification images show colocalization with PSD95 (magenta), apposed to VGluT1 (green) or VGluT2 (green) puncta. Scale bar 500 µm, inset 1 µm.

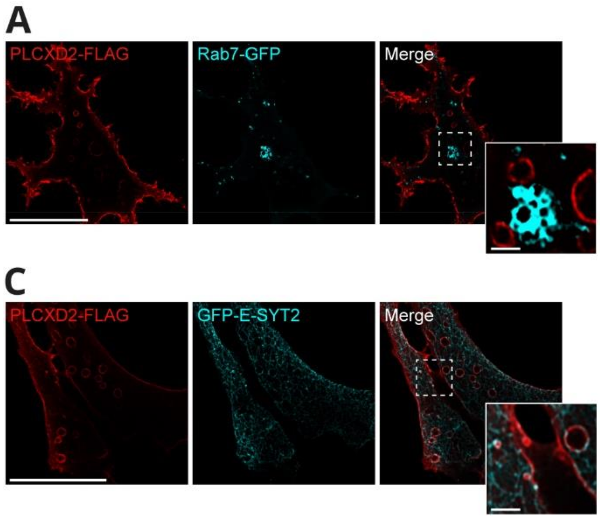


**Supplementary figure 3. GPR158 prevents PLCXD2-induced ER-PM contact site remodeling (See also Figure 2).**

**(A)** HEK239T cells co-expressing PLCXD2-FLAG and Rab7-GFP to label late endosomes were immunostained for FLAG (red) and GFP (cyan). Scale bar 20 µm, inset 2 µm.

**(B)** HEK239T cells co-expressing PLCXD2-FLAG and Rab11-GFP to label recycling endosomes were immunostained for FLAG (red) and GFP (cyan). Scale bar 20 µm, inset 2 µm.

**(C)** HEK239T cells co-expressing PLCXD2-FLAG and GFP-ESYT2 to label ER-PM contact sites were immunostained for FLAG (red) and GFP (cyan). Scale bar 20 µm, inset 2 µm.

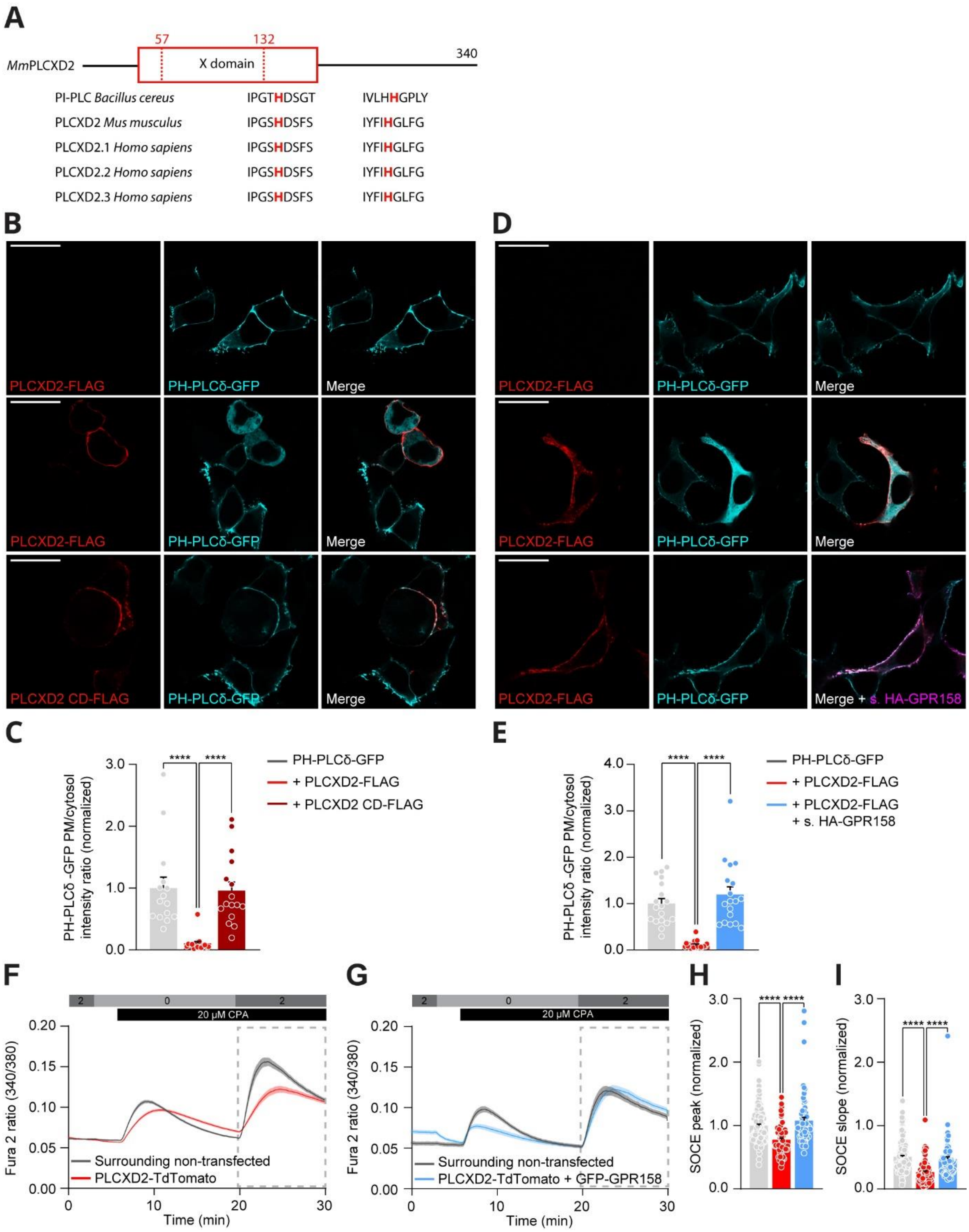


**Supplementary figure 4. GPR158 prevents PLCXD2-induced ER-PM contact site remodeling (See also Figure 2).**

**(A)** Alignment of the amino acid sequences surrounding two conserved histidine residues in the X-domain of PLCXD2.

**(B)** HEK239T cells co-expressing the $PIP_2$ probe PH-PLCδ-GFP together with PLCXD2-FLAG or PLCXD2-FLAG CD immunostained for FLAG (red) and GFP (cyan). Scale bar 20 µm.

**(C)** Quantification of the PH-PLCδ-GFP PM/cytosol ratio. PH-PLCδ-GFP (*n* = 3 experiments, 15 cells), PH-PLCδ-GFP + PLCXD2-FLAG (*n* = 3, 16 cells), PH-PLCδ-GFP + PLCXD2-FLAG CD (*n* = 3 experiments, 15 cells). ****$P$<0.0001, Kruskal-Wallis test, Dunn's multiple comparisons.

**(D)** HEK293T cells co-expressing the $PIP_2$ probe PH-PLCδ-GFP together with PLCXD2-FLAG or together with PLCXD2-FLAG and HA-GPR158 were live labeled for HA (magenta), then fixed, permeabilized and immunostained for FLAG (red) and GFP (cyan). Scale bar 20 µm.

**(E)** Quantification of the PH-PLCδ-GFP PM/cytosol ratio. PH-PLCδ-GFP (*n* = 3 experiments, 18 cells), PH-PLCδ-GFP + PLCXD2-FLAG (*n* = 3, 20 cells), PH-PLCδ-GFP + PLCXD2-FLAG + HA-GPR158 (*n* = 3 experiments, 18 cells). ****$P$<0.0001, Kruskal-Wallis test, Dunn's multiple comparisons.

**(F and G)** HEK239T cells transfected with PLCXD2-TdTomato (red trace) alone or co-transfected with GFP-GPR158 (cyan trace) were loaded with Fura-2 to measure cytosolic Ca2+ levels. Stores were depleted using the SERCA inhibitor CPA (20 µM, black rectangle) in the absence of extracellular Ca2+ (0 mM, light gray rectangle). The SOCE response (dotted area) was then measured following readdition of extracellular Ca2+ (2 mM, dark gray rectangle).

**(H)** Quantification of the peak SOCE response in PLCXD2-TdTomato expressing HEK239T cells (*n* = 4 experiments, 82 cells), HEK239T cells co-expressing PLCXD2-TdTomato and GFP-GPR158 (*n* = 4 experiments, 68 cells), and non-transfected control cells (*n* = 4 experiments, 89 cells). ****$P$<0.0001, Kruskal-Wallis test, Dunn's multiple comparisons.

**(I)** Quantification of the rate of SOCE response in PLCXD2-TdTomato expressing HEK239T cells (*n* = 4 experiments, 82 cells), HEK239T cells co-expressing PLCXD2-TdTomato and GFP-GPR158 (*n* = 4 experiments, 68 cells), and non-transfected control cells (*n* = 4 experiments, 89 cells). ****$P$<0.0001, Kruskal-Wallis test, Dunn's multiple comparisons.

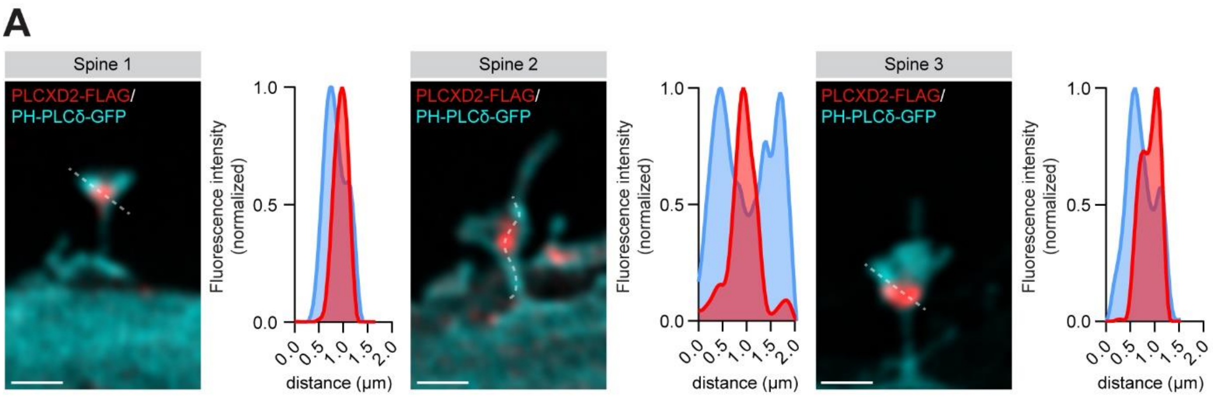


**Supplementary figure 5. PLCXD2 hinders SA formation (See also Figure 3).**

Representative images of dendritic spines from DIV14 cultured mouse hippocampal neurons co-expressing PH-PLCδ-GFP and PLCXD2-FLAG. Fluorescence intensity profile line scans through the spine demonstrate absence of $PIP_2$ at PLCXD2-labeled domains. Scale bar 1 µm.

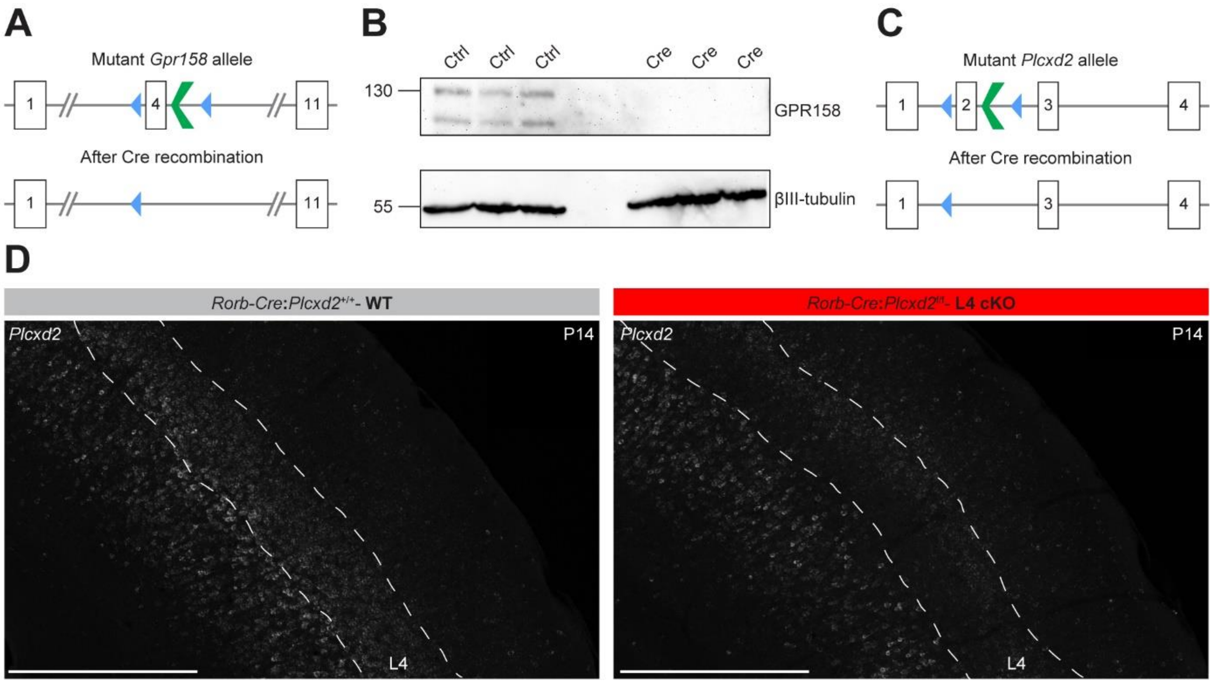


**Supplementary figure 6. GPR158 and PLCXD2 oppositely regulate dendritic spine maturation (see also Figure 4).**

**(A)** Exon 4 of *Gpr158* was floxed to generate a cKO allele following Cre-mediated recombination.

**(B)** Lysates were prepared at DIV14 from cultured *Gpr158*$^{f/f}$ mouse cortical neurons expressing Cre and immunoblotted for GPR158 and βIII-tubulin. Neurons were infected at DIV2 with lentiviral vectors harboring mCherry (control) or Cre-T2A-mCherry (Cre). No GPR158 is detected following Cre-mediated recombination.

**(C)** Exon 2 of *Plcxd2* was floxed to generate a cKO allele following Cre-mediated recombination.

**(D)** P14 mouse coronal brain sections from WT (*Rorb-Cre:Plcxd2*$^{+/+}$) and *Plcxd2* L4 cKO (*Rorb-Cre:Plcxd2*$^{fl/fl}$) mice were probed for *Plcxd2* (gray) using RNAScope at P14. *Plcxd2* expression is strongly reduced in L4 following Cre-mediated recombination. Scale bar 500 µm.

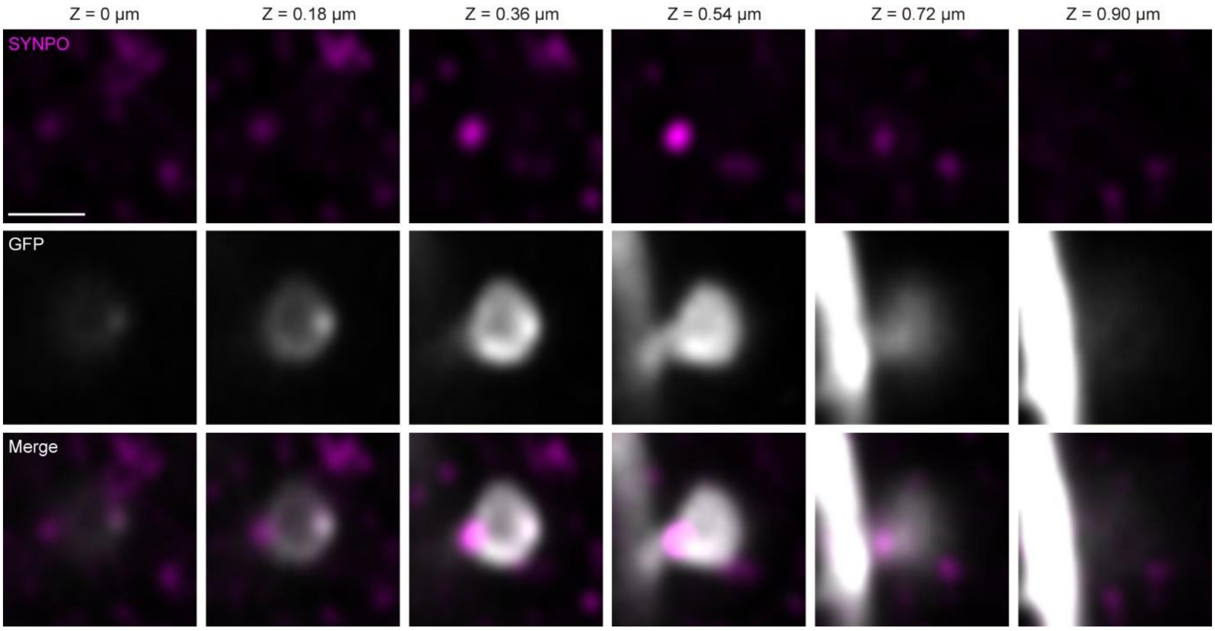


**Supplementary figure 7. A GPR158-PLCXD2 complex regulates SA content and maturation of dendritic spines (see also Figure 5).**

A series of high magnification images at consecutive Z-planes of dendritic spines from *Gpr158*$^{+/+}$:*Plcxd2*$^{+/+}$ mice injected with AAV-TRE-Cre and AAV-SYN-DIO-GFP-IRES-tTA, immunostained for GFP (gray) and SYNPO (magenta) at P14. A spine was considered SYNPO+ when a SYNPO punctum overlapped with the spine head and/or neck at their corresponding Z-planes. Scale bar 1 µm.

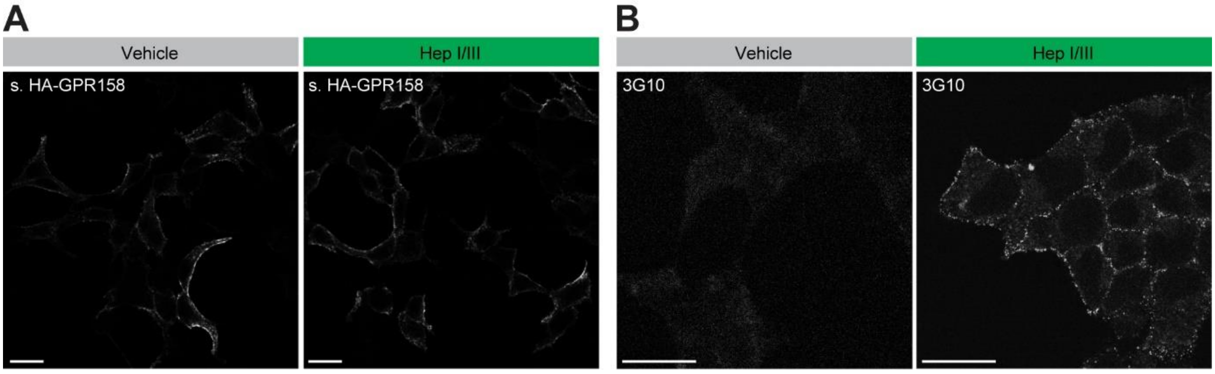


**Supplementary figure 8. Extracellular HSPG binding modulates the GPR158-PLCXD2 complex interaction (see also Figure 6).**

**(A)** HEK293T cells expressing HA-GPR158 treated with vehicle or heparinase I/III for 2 hours were live labeled for HA (gray) to detect the surface pool of GPR158. Scale bar 20 µm.

**(B)** HEK293T cells treated with vehicle or heparinase I/III for 2 hours were immunostained for HS using a 3G10 antibody (gray). Heparinase I/III digestion unmasks an epitope that becomes available for binding of the 3G10 antibody. Scale bar 20 µm.

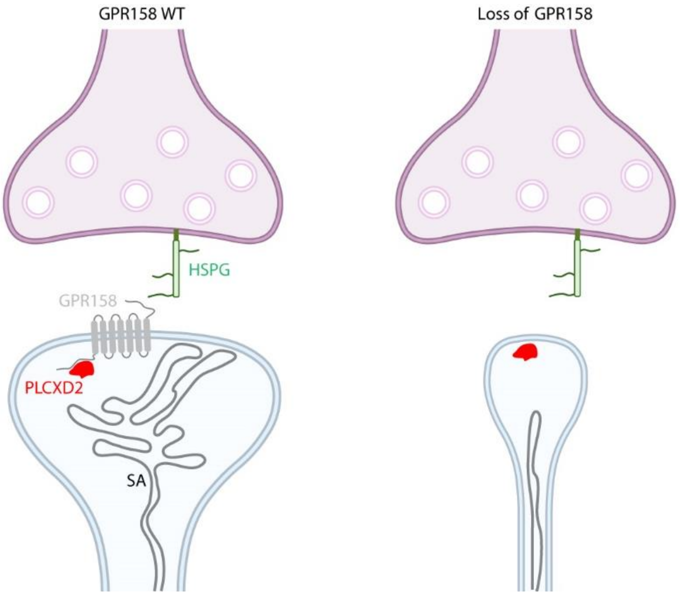


**Supplementary figure 9. Working model.**

Postsynaptic GPR158 limits the activity of PLCXD2 and prevents continuous $PIP_2$ depletion to allow for the formation of a SA and for dendritic spine maturation to occur. In the absence of GPR158, uncontrolled PLCXD2 creates an unfavorable environment that hinders SA incorporation and blunts dendritic spine maturation.
